# InterScale reveals multi-scale cellular interaction programs in spatial transcriptomics

**DOI:** 10.64898/2026.05.07.723456

**Authors:** Francesca Drummer, Sara Jimènez, Federico Di Marco, Anna C. Schaar, Tancredi Massimo Pentimalli, Julia L. Beckman, Nikolaus Rajewsky, Fabian J. Theis

**Author notes:** These authors contributed equally: Francesca Drummer, Sara Jimènez.

## Abstract

Tissue homeostasis and disease emerge from cell–cell interactions operating across spatial scales: from autocrine and juxtacrine signals within micrometers to paracrine gradients coordinating responses across tissues. While these can be read out from spatial transcriptomics, existing computational methods capture either local adjacency-based or long-range dependencies, but rarely both within a single framework. We introduce InterScale, a graph-transformer approach that jointly models local and global cellular interactions from spatial transcriptomics data. By integrating a Graph Convolutional Network as a local component with a global transformer encoder, InterScale learns multi-scale representations of cellular communication. A downstream workflow enables scale-resolved interpretation of interactions from gene to tissue level. Applied to Sonic Hedgehog morphogen patterning in neural organoids, InterScale resolves spatially restricted neuronal differentiation programs and broader progenitor regulatory states along the morphogen gradient. In a human pancreatic dataset contrasting healthy and type 1 diabetic tissue, it reveals disease-associated spatial reorganization and tissue remodeling. InterScale’s modular architecture supports diverse spatial transcriptomics platforms and provides a scalable, unbiased, and biologically interpretable framework for studying cellular interactions across scales.

## Introduction

Cell–cell interactions are essential for maintaining tissue homeostasis, coordinating developmental processes, and responding to environmental changes^1^. These interactions occur across multiple spatial scales, from signals acting on the same cell to those coordinating communication across entire tissues^2^ (Suppl. Fig. 1A). At the smallest scale, autocrine signaling occurs when cells respond to signaling molecules they secrete themselves, enabling self-regulation of cellular states. Juxtacrine signaling operates through direct physical contact between neighboring cells via membrane-bound ligands and receptors, highlighting the importance of spatial proximity in specific signaling pathways. At slightly larger spatial ranges, paracrine signaling involves the release of diffusible molecules that act on nearby cells within the local microenvironment. Finally, endocrine signaling represents long-range communication, where hormones travel through the bloodstream to regulate distant target tissues^3^. These multiscale communication mechanisms are fundamental to the spatial organization of tissues. Gradients of morphogens and other diffusible signaling molecules can propagate information across tissue space, coordinating cell fate decisions during development^4,5^ and driving tissue remodeling in physiological and pathological contexts. Disruption of these multiscale cell–cell interactions are often involved in diseases, including cancer^6^ and autoimmune disorders such as type 1 diabetes^7^, where altered signaling pathways result in abnormal cellular behavior.

Computationally, inferring cell-cell communication (CCC) across different length scales still remains a challenge, due to data limitation and missing ground truth^8^. From the roughly fifty CCC methods that incorporate spatial proximity^9,10^ (Suppl. Table 1), existing methods mostly model cell-cell communication at a single spatial scale, either local neighborhood interactions or broader tissue context, but do not integrate both within a unified framework.. Graph-based and neighborhood-aware models capture local effects, but typically operate within fixed radii or adjacency-defined neighborhoods, remaining inherently short-range which limits their ability to detect distal or gradient-driven interactions. Methods such as NCEM^11^ models niche effects using cell-type labels or NicheCompass^12^ rely on ligand–receptor databases^13,14^, provide valuable biological priors but are incomplete, and biased toward known pathways and more infeasible for imaging-based spatial transcriptomics technologies with limited gene space. Others, like MISTy^15^, distinguish local from broader tissue signals but rely on predefined spatial kernels that tend to down-weight long-range contributions, follow up methods like Kasumi^16^, aggregate expression values of local neighborhoods across within a patch, losing fine grained signalling patterning. Attention-based models, such as STEAMBOAT^17^ and AMICI^18^ demonstrate the power of learned interaction mechanisms, yet they still do not provide a unified representation that simultaneously captures local and global spatial influences while remaining unbiased with regard to pathway annotations or imposed distance scales. As a result, current tools address CCC by modeling either local adjacency-based communication or generic long-range dependencies, but rarely both in an integrated and unbiased framework.

Hence, despite these approaches, a key challenge remains, namely how to represent cellular interactions across multiple spatial scales. Existing machine learning architectures capture different aspects of this problem. Graph neural networks (GNNs) naturally encode local spatial structure but are constrained by fixed receptive fields and therefore struggle to capture long-range dependencies^19^graph. Conversely, Transformers^20^ excel at capturing global relationships but lack inductive biases for spatial structure and can over-attend to distant cells. A framework capable of integrating these complementary scales of information would provide a more complete representation of cellular interactions within tissues.

Here we present InterScale, a modular framework for modeling cellular interactions across spatial scales in spatial transcriptomics data. InterScale integrates graph-based representations of local neighborhoods with transformer-based attention to capture both short-range interactions and broader tissue-level context within a unified architecture. This approach decomposes each cell’s transcriptional state into contributions from intrinsic programs, neighboring cells, and distal signals across the tissue. To interpret these representations, we introduce a gene evaluation pipeline based on standardized decoder loadings that identifies scale-specific driver genes in a cell-type-agnostic manner. By avoiding reliance on ligand–receptor priors or predefined cell-type labels, InterScale enables unbiased discovery of multiscale communication patterns, revealing both localized cellular niches and tissue-wide signaling programs.

## Results

### InterScale models local and global interactions in spatial transcriptomics

InterScale is designed to integrate local interactions (e.i. direct signalling) in a global tissue context (Suppl. Fig. 1A). The model learns interaction patterns directly from spatial omics data consisting of one or multiple samples from the same tissue with one or more conditions. We represent every sample as a graph, where each cell or spot *N* is a node with gene expression features *F* and edges encode spatial proximity (Suppl. Fig. 1B). Each sample is described by a gene expression matrix *X* and an adjacency matrix *A*, and can be processed either as a whole or subdivided into sliding windows for scalability (Suppl. Fig 1B, methods and Suppl. methods). During training, InterScale employs a self-supervised learning strategy based on masked nodes^21^, supporting both classification (graph- or node-level label prediction) and regression (gene expression reconstruction) tasks.

To achieve the joint learning of local and global interactions, InterScale builds on the GraphTrans architecture^22^ (see Suppl. methods for architectural differences): a Graph Convolutional Network (GCN)^23^as local modelling component and a multi-head self-attention transformer^20^ as global modelling component (Fig. 1A). The local component propagates information only from *k*-hop connected neighbouring nodes, where *k* corresponds to the number of GCN layers (Fig. 1B). Hence, the local component creates a locally informed neighborhood embedding for each cell which we will refer to as local embedding *H*_*local*_. Next, the transformer takes the locally informed embeddings and an additional classification token (CLS) that represents the entire tissue context by being connected to each cell, enabling tissue or sliding window downstream tasks. The global interactions arise from the fact that a transformer encoder without additional positional embeddings or encodings assumes a fully connected graph between the embeddings (Fig 1B). The output of the transformer will be referred to as global embedding *H*_*global*_, as it captures long-range interactions across the entire tissue or sliding window space. Separate linear decoder for each embedding learns to predict the reconstructed gene expression *X*. The modularity of InterScale’s architecture allows the local and global components in the model to be easily adjusted by exchanging it with either a different GNN architecture, such as graph-isomorphism network^19^ or a spatial domain aware embedding from other models in the spatial transcriptomics domain (i.e. CellCharter^24^, BANKSY^25^). These models can either be trained within the InterScale architecture or their trained outputs loaded as precomputed embeddings to replace a trainable local component (Suppl. Fig. 2).

**Figure 1:**
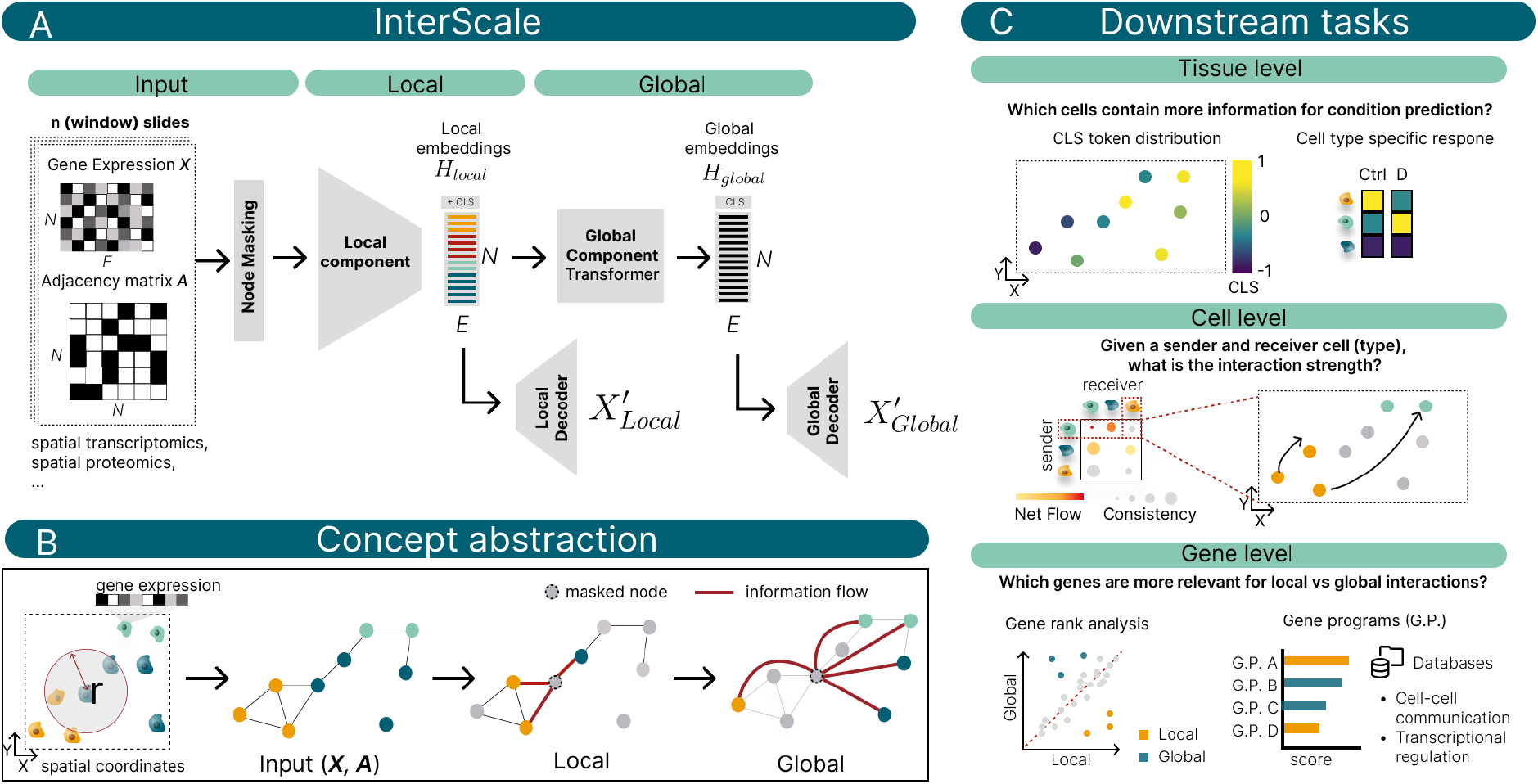
Overview of InterScale. A) InterScale takes multi-sample spatial omics data with cell- or spot-level observations as input. The features are represented as gene expression matrix and spatial connectivities as adjacency matrix to the model. A fraction of nodes are masked and passed to the Local Component (e.i. GCN). The Local Component encodes each cell as an embedding with the local embeddings representing the local connections of each cell. The local embeddings and additional classification token (CLS) are passed to the Global Components (e.i. self-attention transformer) summarizing the global interactions of cells in the global embedding. Specific for each component the reconstruction of the gene expression matrix is computed using the Local and Global Decoder. B) Abstracting the workflow of InterScale concept, one can consider a connected graph where the masked node is predicted using information from i) local information considering only connected nodes and ii) global information considering all remaining nodes. C) Multi-scale downstream tasks are on i) tissue level, ii) cell level and gene level. Tissue level uses the CLS to summarize cell specific responses about which cell type might contribute more to a condition prediction. Cell level analyses sender and receiver cell interaction strength using the attention matrix and gene level informs about genes that are relevant for local or global signalling.

As a computational toolbox, InterScale provides functions for multi-scale analysis across three scales: tissue, cell and gene level (Fig. 1C). Both graph and cell level-based downstream tasks use the attention matrix, while the gene level evaluation uses the local and global embeddings. For robust evaluation of the attention matrix, we use attention probing to quantify the relevant attention across all layers and heads^26^ (see methods). First, graph level evaluation uses the CLS in the attention matrix to identify which cells contribute to the condition prediction^27^. Second, normalised attention scores at the cell level inform interaction strength between sender and receiver cells. Lastly, at the gene level, analysis of local and global embeddings enables ranking of gene contributions and identification of pathway-driven activity across embedding dimensions.

### Global structures contains relevant tissue information

In classification tasks, short-range spatial context, typically limited to neighborhoods of up to ~50 cells, has often been shown to be redundant^11,28,29^. This is because local neighborhoods primarily reflect immediate transcriptomic similarity or local cell-type composition, failing to add significant discriminative information beyond the single-cell level for predicting labels such as cell type or tissue state. We hypothesized that for robust classification, particularly of complex tissue states, a substantially larger, global tissue context is required. To assess whether contribution of tissue-scale context enhances predictive performance, we systematically evaluated the relative importance of local neighborhood information and global tissue-level structure across two classification tasks: graph-level label prediction and node-level label prediction. To that end, we compared models leveraging only local information (Local Component; GCN), only global information (Global Component; multi-head self-attention transformer), and their combination (InterScale).

For graph label prediction, we predict the condition or disease stage from a given tissue sample. We first compare the class-specific test F1 scores on a 10X Visium Alzheimer’s disease dataset^30^, which includes a total of six slides (three healthy and three mid-stage AD patients). In this setup, every spot is connected to its neighboring spots, forming a hexagonal grid. To increase the sample size, we split the dataset into sliding windows (Fig. 2A), allowing us to predict conditions from windows rather than whole tissue slides. The overall accuracy and class-specific F1 scores for InterScale are higher than those for the GCN (Fig. 2B).

**Figure 2:**
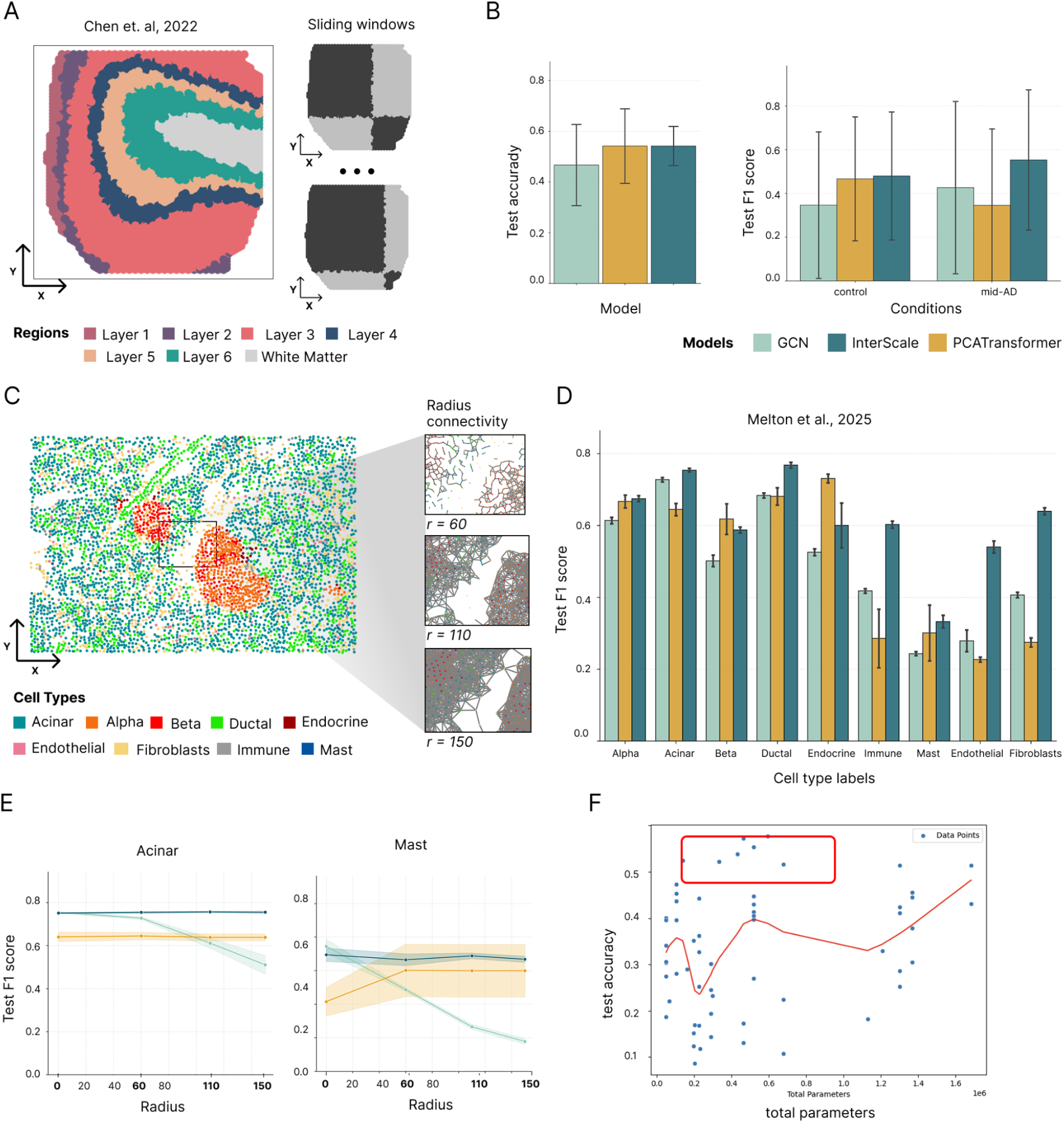
InterScales prediction performance improves with tissue level information and demonstrates multi-scale robustness across diverse spatial transcriptomics platforms. A-B: human brain Visium dataset from Chen et al., 2022 (N=6, number of sliding windows = 46) with three control and three mid Alzheimer disease (AD) samples from the middle temporal gyrus (MTG). A) Representative spatial slide from Chen et al., 2022 colored by anatomical layers and the alternating sliding window grid (black and white squares) used as input to the model. B) Comparison of model performance in classifying disease states (Control vs. Mid-AD). Bar-and-whisker plots display the overall test F1-score (left) and condition-specific F1-scores (right) for three architectures: GCN (local component, light blue), InterScale (local & global model, dark blue), and PCA-Transformer (global component, yellow). C-F: human pancreas CosMx dataset (Melton et al., 2025) (N=72, sliding_windows = 596) C) Representative CosMx slide colored by cell-type identity. Zoomed-in panels illustrate the resulting graph topology when varying the neighborhood search radius (r), ranging from sparse to dense connectivity. D) Predictive performance (test F1-score) across distinct cell types across three model architectures (colors). E) Model stability across varying graph radii (r) for two cell types, mast and F) Parameter space against cell type prediction performance, each dot indicating a model with specific parameter settings and parameter space.

However, both models are susceptible to overfitting on either condition, resulting in a high standard deviation across conditions. We hypothesize that this high variance arises from the limited patient samples, as each training, test, and validation split only sees one representative patient per condition. Observing the condition-specific test F1 score performance of the models on three different datasets confirms this hypothesis (Suppl. Fig. 3A). Across all three models, we observe a decrease in generalizability with higher patient-to-patient variability and lower samples per patient (Suppl. Fig. 3A). In line with previous studies^31–33^, global information models (PCATransformer and InterScale) are less affected by condition invariance. This suggests that tissue-level information represents a different source of information compared to neighborhood information. Even without major improvements to InterScale’s ability to better capture global tissue patterns, we demonstrate that the CLS token from the global component provides an intuitive way to analyze whether some cells contribute more to the classification of one cell type than others (Section 4).

In addition to graph label classification, we tested the selected models for node label classification. We use a CosMx Type 1 Diabetes Pancreas dataset. Compared to spot-based datasets, such as Visium from above, cellular resolution datasets are usually modelled using kNN, delaunay triangulation or radius-based graphs. While there is no common consensus on which representation is most meaningful, we use a radius-based representation (Fig. 2C). For radius-based neighborhoods we observe an indirect relation between node connectivity and density changes across the tissue. In line with previous work, cell type classification performance across models is relatively consistent (Fig. 2D) supporting that transcriptomics differences alone often drive prediction tasks for cell types and that spatial information might be redundant. However, similar to other graph transformer models^31,34^, we observe that classification performance improves with local and global context for abundant cell types in unbalanced datasets such as immune and endothelial cells. Taking this one step further, to a spatially informed label prediction such as brain region classification, we observe a better performance for InterScale that has global information available (Suppl. Fig. 3C). We hypothesise that global information is valuable for highly affected regions, such as Layer 2, 3 and 4 that show preferential pathology accumulation leading to varying proportion and gene count distribution of spots in AD^35^. Finally, a common observation for radius-based neighborhood selection approaches is the sensitivity to radius selection for specific cell types, as one assumes that cell types communicate across different ranges^11^ or different topological information are important for different cell types^36^. By including global context we reduce the sensitivity to radius selection (Fig. 2E). Again, we confirm that for some cell types, such as Acinar cells, spatial information and their connections to other cell types is more predictive than for others (e.g. Alpha, Ductal - see Suppl. Fig 3C).

Beyond performance-based metrics, we evaluated the trade-off between increasing model complexity (higher parameter space architectures) and performance. In line with other publications^31^, we found that model performance often stagnates or even reduces as the number of parameters increases. The best-performing models were consistently found within a lower parameter space. This stagnation with increasing parameters has been previously observed in Graph Convolutional Networks (GCNs) due to oversmoothing, but our findings indicate it also holds true for Transformer architectures. We find specific model parameters, such as the embedding dimension *E* have more influence on the performance than others such as the GCN number of layer *L*_*GCN*_.

We demonstrated that the predictive performance improves when incorporating the global component (Fig. 2B,D,E), providing direct evidence that tissue-scale structure contains biologically relevant information. To further analyse neighborhood level interactions, we went on to focus on gene expression prediction instead of classification hereafter.

### The local and global role of the SHH morphogen along the dorsoventral axis

To demonstrate the ability of InterScale to distinguish local and global transcriptional programs, we analyzed a spatial transcriptomics dataset profiled using molecular cartography, where Sonic Hedgehog (SHH) signaling was optogenetically induced to drive dorsoventral patterning in neural tube organoids (Fig. 3A)^37^. This system provides biological proof of concept for the type of multiscale spatial organization that InterScale is designed to detect, as morphogen-driven patterning generates transcriptional programs that vary across both local and tissue-scale gradients. The dataset consists of four control and four SHH-induced organoids, in which graded activation of the SHH pathway establishes spatial domains corresponding to ventral and dorsal neural identities. Consistent with this perturbation, SHH induction results in the emergence of ventral progenitor markers, including *NKX2-2, NKX6-1, PAX6, OLIG1/2, IRX3*, and *DBX1/2* (Fig. 3B). To characterize the length scale at which these genes are organized, we quantified spatial autocorrelation using Moran’s I^38^ across progressively larger kNN neighborhoods. Genes whose Moran’s I rapidly decreases as the neighborhood size *k* increases correspond to spatially localized patterns, whereas genes with a more gradual decay exhibit broader, tissue-scale organization (Fig. 3C). Hence, the dependence of Moran’s I on neighborhood size can be interpreted as an effective spatial length scale. Focusing on gene-level spatial organization, we demonstrate that InterScale can identify spatial programs in a cell–type–agnostic manner without relying on predefined cell-type annotations.

**Figure 3:**
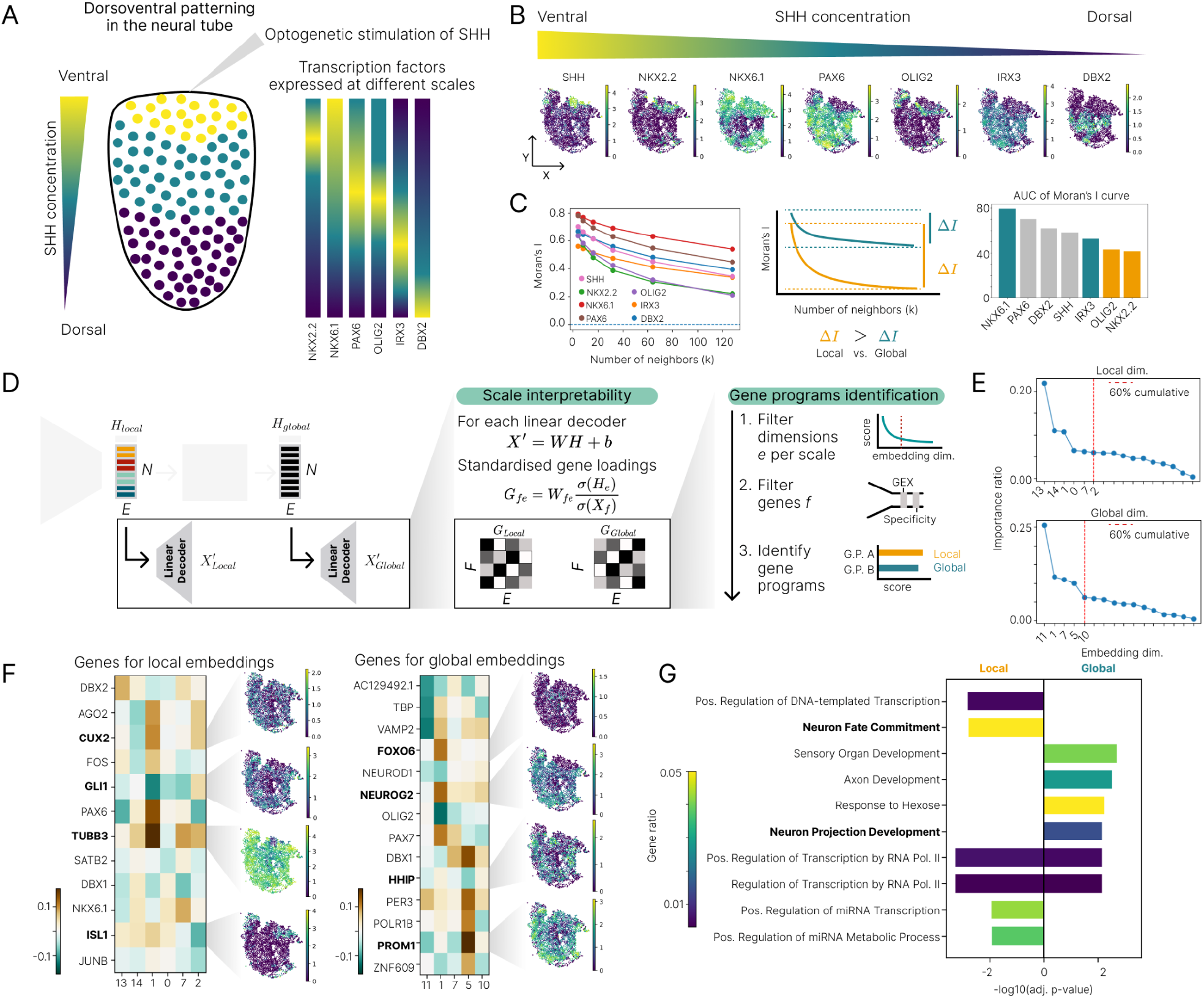
InterScale identifies transcriptional programs operating at distinct spatial length scales during SHH-driven dorsoventral patterning. A) Schematic of the experimental system. Optogenetic stimulation of Sonic Hedgehog (SHH) signaling in neural tube organoids induces dorsoventral patterning, generating transcriptional programs that vary across spatial scales along the SHH morphogen gradient. B) Spatial expression of representative ventral neural progenitor markers across the tissue. C) Moran’s I was computed across increasing neighborhood sizes to quantify spatial autocorrelation at different spatial scales. The area under Moran’s I curve summarizes the spatial scale of each gene, where rapidly decreasing autocorrelation corresponds to more localized patterns, while gradual decay indicates broader spatial organization. D) Overview of the InterScale interpretability framework. Local and global embeddings are decoded through linear decoders to reconstruct gene expression. Decoder weights are transformed into standardized gene loadings, which quantify the change in gene expression (in gene standard deviation units) associated with a one–standard deviation change in an embedding dimension. These loadings enable identification of gene programs associated with local and global spatial embeddings. E) Selection of informative embedding dimensions. Embedding dimensions are ranked according to their contribution to gene expression variability, and dimensions explaining the majority of the cumulative variance are retained separately for local and global embeddings. F) Genes associated with selected embedding dimensions. Heatmaps show standardized gene loadings for representative genes across selected local and global embedding dimensions. Corresponding spatial expression maps illustrate the spatial patterns captured by each embedding dimension. G) Functional enrichment analysis of gene programs derived from local and global embeddings. Local programs are enriched for neuronal differentiation and morphogen-response pathways, whereas global programs are enriched for broader developmental and transcriptional regulatory processes

To verify whether inferred local and global embeddings capture meaningful and distinct spatial information, we first assessed their structure and relationships. Given the sequential nature of InterScale’s architecture, we tested whether the two embeddings capture different signals rather than converging toward similar representations. Clustering of the embeddings revealed coherent spatial patterns corresponding to distinct transcriptional domains (Suppl. Fig. 5A). In addition, correlation analysis across embedding dimensions within each length scale showed low pairwise correlations (Suppl. Fig. 5B), indicating that the learned components are largely linearly independent. Together, these analyses indicate that the embeddings encode multi-scale spatial information and represent largely linearly independent features of the data. Next, to identify the transcriptional programs captured by these embeddings, we leveraged the interpretability of the linear decoders used to reconstruct gene expression from the latent representations (Fig. 3D). In this framework, the decoder weights quantify the contribution of each embedding dimension to the expression of individual genes. To make these contributions comparable across genes and embedding dimensions, we computed standardized gene loadings by scaling the decoder weights by the ratio of the variability of the embedding dimension and the gene across cells. This normalization provides a scale-independent measure of the influence of each latent dimension on gene expression. Using these loadings, we implemented a gene program identification pipeline separately for the local and global embeddings (Methods). First, we selected the most informative embedding dimensions *e* at each spatial scale based on their overall contribution to gene expression variability. Second, we identified genes specific to individual embedding dimensions by prioritizing genes with strong standardized loadings while favoring highly variable genes to focus on biologically informative signals. Finally, the resulting gene sets derived from local and global embeddings were associated with biological programs through functional enrichment analysis, enabling the identification of transcriptional programs operating at distinct spatial length scales.

Using the pipeline described above, we next identified the most informative embedding dimensions for each spatial scale (Fig. 3E). For the selected dimensions, genes were ranked according to their standardized loadings, and the union of the top 10 genes per embedding dimension was visualized to highlight the transcriptional programs captured by the model (Fig. 3F). Functional enrichment analysis of these gene sets revealed distinct biological programs associated with the local and global embeddings (Fig. 3G). The gene programs associated with the local embeddings included markers of neuronal differentiation and transcriptional responses to morphogen signaling. In particular, *GLI1*, a transcriptional effector of the Sonic Hedgehog (SHH) pathway, and ISL1, a motor neuron differentiation factor, link these programs to ventral neuronal specification downstream of SHH signaling^39^. The presence of *TUBB3*, a marker of neuronal maturation, and *CUX2*, involved in cortical neuron specification, further suggests that local embeddings capture spatially restricted neuronal differentiation states^40^ (Fig. 3F-G). In contrast, gene programs associated with the global embeddings were enriched for regulators of progenitor states and developmental signaling. These included *PROM1*, a neural stem cell marker^41^, *NEUROG2*, a proneural transcription factor controlling early neurogenesis^42^, and HHIP, a negative regulator of SHH signaling that modulates morphogen gradients^43^, suggesting that global embeddings capture broader developmental and regulatory programs operating across the tissue (Fig. 3F-G).

To further validate these findings, we performed a complementary analysis comparing gene reconstruction across the local and global decoders (Methods). Because each decoder preferentially reconstructs genes operating within its effective spatial scale, we ranked genes according to their reconstruction performance for each decoder and examined the difference in rank between the two. Genes well captured by the local decoder therefore exhibit high ranks in the local model but low ranks in the global model, and vice versa. This analysis confirmed that genes previously identified in our embedding analysis, such as *ISL1* and *FOXO6*, preferentially associate with local and global spatial programs, respectively (Suppl. Fig. 5C).

Taken together, InterScale accurately recovers transcriptional programs operating at distinct spatial length scales. By separating local and global embeddings, the framework resolves spatially restricted differentiation signals from broader developmental regulatory programs within the same tissue. In the context of SHH-driven dorsoventral patterning, InterScale captures both localized neuronal specification programs and global progenitor and morphogen-regulated states. Finally, these signatures were identified directly from gene-level spatial structure in a cell-type-agnostic manner, highlighting InterScale’s ability to uncover biologically meaningful multi-scale spatial organization without prior information about cell type annotations.

### Global tissue context reveals condition-specific spatial programs in the human pancreas

To evaluate whether InterScale can extract biologically meaningful spatial programs from more complex datasets, we next applied it to a human pancreas spatial transcriptomics dataset generated using the image-based platform CosMx^7^ (Fig. 4A). In contrast to the controlled morphogen-driven system analyzed above, this dataset captures native tissue organization and includes both non-diabetic (ND) and type 1 diabetic (T1D) samples, introducing additional sources of variability such as donor heterogeneity and disease-associated remodeling. The pancreas contains well-defined spatial niches, most prominently the endocrine islets embedded within the surrounding exocrine tissue, alongside diverse stromal and immune populations. These structural features represent a challenging but biologically relevant setting to assess whether InterScale can disentangle transcriptional programs across multiple spatial scales while remaining robust to differences in tissue architecture and condition-specific changes.

**Figure 4:**
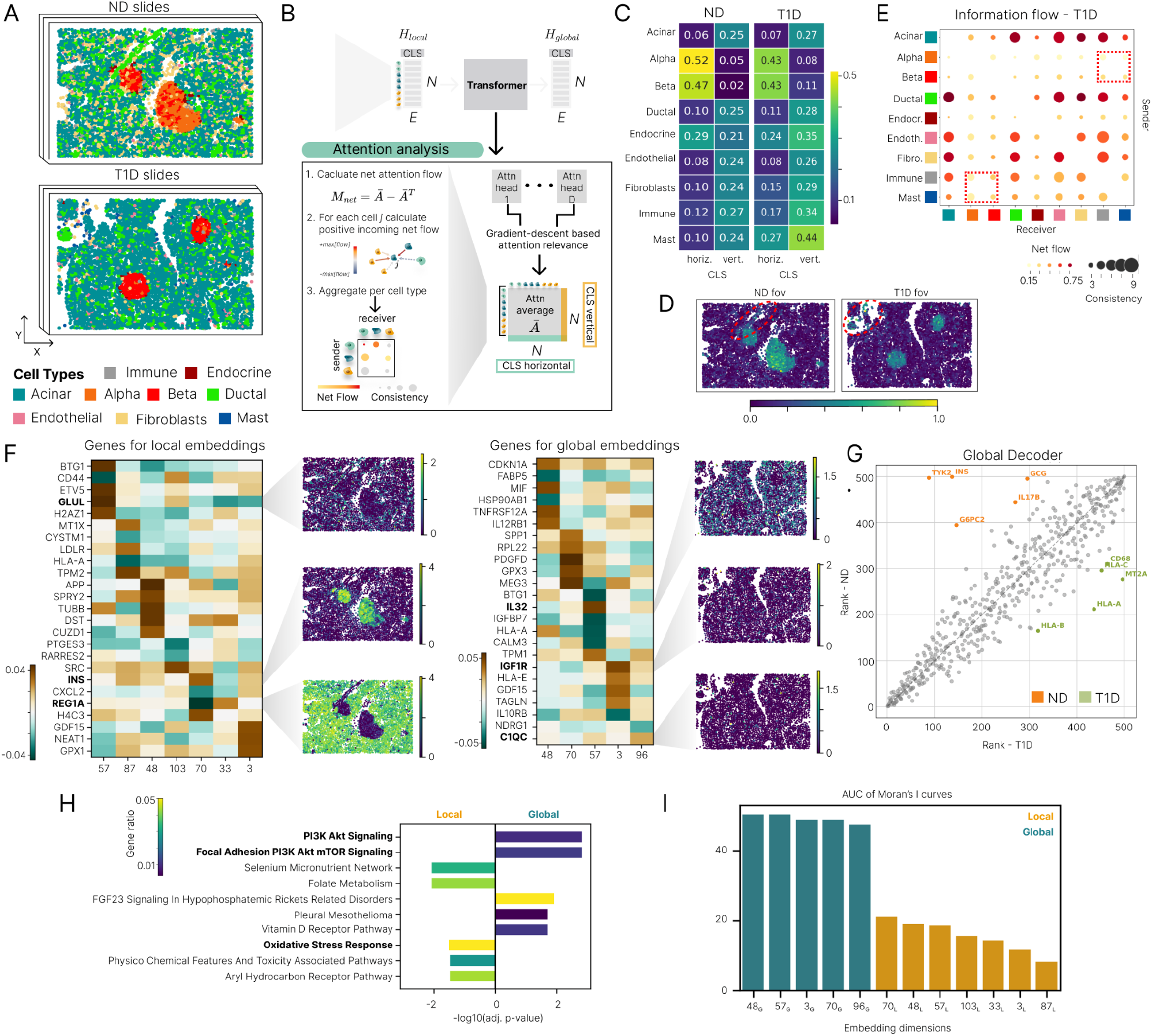
InterScale identifies multiscale interaction programs in human pancreas spatial transcriptomics. A) Spatial organization of representative non-diabetic (ND) and type 1 diabetic (T1D) CosMx slides colored by annotated cell types. Endocrine islets are embedded within the surrounding exocrine and stromal tissue. B) Overview of the attention-based interaction analysis. The transformer processes local and global embeddings, augmented with a CLS token that represents tissue context. Attention heads are aggregated using gradient-based relevance scores to obtain a unified attention matrix. Net attention flow is computed by subtracting incoming and outgoing attention, and interaction scores are summarized across cell types to estimate sender–receiver relationships and interaction consistency. C) Cell-type contributions to the CLS token for ND and T1D conditions. Horizontal and vertical CLS attention scores represent receiver and sender relevance respectively, highlighting the contribution of different cell types to tissue-level representations. D) Spatial distribution of CLS-derived relevance scores projected onto the tissue for ND and T1D slides, illustrating condition-associated regions contributing to classification. E) Sender–receiver information flow map inferred from the attention matrix for T1D samples. Dot size represents interaction consistency across slides and color indicates net attention flow, highlighting bidirectional interactions between endocrine and immune populations. F) Genes associated with local and global embeddings derived from standardized decoder loadings. Heatmaps show top-ranked genes across informative embedding dimensions, with example spatial expression patterns illustrating localized versus tissue-scale transcriptional programs. Local genes spatial expression are shown in ND slide while global gene expression are shown in T1D slide. G) Comparison of global decoder gene rankings between ND and T1D conditions. Endocrine-associated genes are preferentially ranked in ND, whereas immune-related genes show higher ranks in T1D. H) Functional enrichment of gene programs derived from local and global embeddings. Local programs are enriched for cell-state-specific processes such as oxidative stress responses, whereas global programs highlight intercellular signaling pathways including PI3K–AKT signaling and focal adhesion. I) Spatial autocorrelation analysis of representative embedding dimensions using Moran’s I. The area under the moran’s I curves across increasing neighborhood sizes shows that global dimensions maintain higher autocorrelation across larger spatial scales resulting in higher AUCs, while local embedding dimensions exhibit rapid decay of spatial autocorrelation, resulting in lower AUCs.

We investigate how InterScale captures spatial context in this complex tissue by leveraging the CLS token as a proxy of tissue level interactions (Methods). To interpret the interactions, we extract *D* attention heads across *L* transformer layers and employ a gradient-based attention relevance approach (Methods) to collapse these multi-head signals into a single, unified attention map 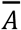 (Fig. 4B). From the attention matrix 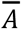 we obtain vertical and horizontal scores for the CLS token which can be interpreted as “from cell type” (sender) and “to cell type” (receiver) scores respectively^27^. The normalized CLS token (Methods) can be interpreted by cell type specific relevance for a given condition (Fig. 4C) or spatial distribution of condition classification (Fig. 4D). According to classification token, the most relevant cell types for both control and T1D prediction are Alpha and Beta cells in the islets (Fig. 4C). This might stem from the fact that both cell types are most variable across conditions. Furthermore, we observe that Mast cells exhibit a stronger signal in T1D compared to ND, consistent with previous studies that have found mast cells infiltrate pancreatic islets in samples from donors with T1D^44^.

While the CLS token summarizes tissue-level interaction patterns, a more granular view of cellular communication can be obtained directly from the attention matrix 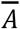 encoding directional interactions between individual cells (sender-receiver relationships)^17,18,45^. To evaluate whether these attention scores capture meaningful interaction patterns, we compared InterScale’s attention matrix with those produced by AMICI and Steamboat (Suppl. Fig. 4A). AMICI showed the highest correlation with Steamboat’s local attention matrix, suggesting a focus on local interactions, whereas InterScale correlated with both local and global Steamboat matrices. Additional comparisons using the Legnini dataset revealed that most Steamboat attention heads capture limited signal, with attention concentrated in the ego head (Suppl. Fig. 4E–F). Consistent with previous observations, correlations among attention matrices remained modest, reflecting the sensitivity of attention magnitudes to the attribution methods used. We therefore focus our interpretation primarily on the directionality of interactions rather than absolute attention magnitudes.

To focus on directionality instead of magnitude of the attention matrix, we aggregate the positive net attention flow *M*_*net*_ per cell type obtaining a sender-receiver information flow map (Fig. 4B). For T1D the sender-receiver information flow map (Fig. 4E) shows bidirectional information exchange between Islet cells (Alpha and Beta) and immune cells (Immune and mast). Next, we compare the CLS token analysis with GraphCompass, a spatial graph analysis method that assesses a pairwise similarity test between ND and T1D (Suppl. Fig. 6A) on fine grained cell types. In contrast to tools that perform quantitative testing on tissue level such as GraphCompass^46^, InterScale’s CLS token has the advantage of being projected onto the spatial slide to highlight regions of interest, such as an isolated region of ductal cells in ND and mast cells in the T1D (Fig. 4D).

While attention-based analyses reveal patterns of cellular communication, the decoder representations allow us to interpret the transcriptional programs underlying these interactions. For the gene evaluation, we leveraged the standardized gene loadings derived from the local and global decoders to rank genes associated with the most informative embedding dimensions (Suppl. Fig 6C). Genes classified as local were associated with cell type specific programs, including INS, REG1A, and GLUL, which mark endocrine and epithelial cell states^7^. In contrast, genes classified as global were dominated by immune and stromal signaling pathways, including *C1QC, IL32*, and *IGF1R*, reflecting inflammatory and paracrine programs that propagate across broader tissue regions^47^ (Fig. 4F). Consistently, comparison of global decoder gene ranks between ND and T1D highlighted condition-specific differences, with endocrine genes ranked higher in ND and immune-associated genes ranked higher in T1D (Fig. 4G). Functional enrichment analysis further highlighted distinct pathway signatures between the two spatial scales (Fig. 4H). Local gene programs were enriched for cell-state-specific processes, including oxidative stress response, consistent with metabolically active epithelial and endocrine niches within the pancreas^48^. Global programs were enriched for signaling pathways linked to intercellular communication and tissue remodeling, such as PI3K–AKT signaling and focal adhesion pathways, which are commonly associated with stromal and immune-mediated microenvironmental interactions^49^. The spatial autocorrelation profiles of the embedding dimensions further supported these differences. When computing Moran’s I across increasing neighborhood sizes, dimensions associated with local embeddings showed a pronounced decay in spatial autocorrelation (Suppl. Fig. 6E), indicative of spatially confined expression patterns. In contrast, global embedding dimensions declined more gradually, consistent with transcriptional programs that extend across larger regions of the tissue. We showed the persistence of spatial structure across scales by computing the area under the Moran curve, features with higher AUC values indicate a more persistent spatial organization (Fig. 4I).

Together, these analyses demonstrate that InterScale captures biologically meaningful spatial programs across multiple scales in complex human tissues. The CLS token highlights tissue-level interaction patterns and condition-specific cellular contributions, while the attention-based sender–receiver analysis reveals directional communication between endocrine and immune populations. Additionally, analysis of the decoder-derived gene programs further distinguishes spatially localized cell-state signatures from broader tissue-scale signaling processes. Collectively, these results indicate that InterScale can disentangle local cellular programs from global tissue-level interactions and reveal disease-associated spatial reorganization in the human pancreas.

## Discussion

InterScale’s key contribution lies in its capacity to jointly learn local and global embeddings, enabling biologically interpretable, multi-scale dissection of cell-cell interactions. Our tool successfully identifies the gene programs that act at the local and global tissue scales during dorsoventral patterning of the neural tube. Moreover, in more complex datasets, including spatial slides from adult pancreas of healthy and T1D individuals, InterScale demonstrated superiority in classifying the conditions, while also providing insights into the sender-receiver flow of information. Furthermore, the identified local and global gene programs accurately described patterns of inflammation occurring at the global tissue scale due to T1D, and metabolic signatures reflected proper local interactions in the islets of Langerhans.

A central design principle of InterScale is the integration of graph-based and transformer-based architectures to jointly model neighborhood and tissue-level context. In most experiments, local interactions are captured using a graph convolutional network, while global relationships are modeled through a multi-head self-attention transformer encoder. Importantly, the framework is modular, enabling the local component to be replaced with alternative GNN variants or spatial domain detection methods, and the global transformer encoder can be substituted with architectures like sparse attention transformers that reduce computational cost^50–52^ or cross-attention transformers^18^ to integrate information across modalities simultaneously or infer directional ligand-receptor interactions (Supp. Fig. 2). We demonstrated the versatility of this modular design by applying InterScale across datasets spanning diverse spatial transcriptomics technologies with varying gene panel sizes and spatial resolutions. Modelling interactions at the level of cellular neighbourhoods, rather than individual cells, may contribute to this robustness, as it mitigates sensitivity to technical artefacts such as gene spillover or segmentation errors. More broadly, our results suggest that spatial transcriptomic signals can be decomposed into distinct components operating at different spatial scales, providing a principled framework for separating local cellular states from tissue-level regulatory context.

In this study classification tasks serve as a quantitative probe to assess whether local and global tissue context contain biologically relevant information. The main contribution of InterScale lies in the joint learning of local and global representations, enabling a richer post hoc interpretation of cellular communication across spatial scales. This highlights that the primary strength of InterScale lies not only in predictive performance, but in its ability to provide interpretable, multi-scale insights into tissue organisation. Future extensions could further enhance condition-specific modelling, for example by explicitly incorporating condition-dependent latent variables or additional integration strategies. However, exploring these directions is beyond the scope of the present study.

The interpretability of these interactions arises from the combined analysis of attention-derived interaction scores and embedding-based gene programs. We found that direct use of raw attention values can be unreliable due to attention sink effects and their dependence on sequence length. Consistent with previous research, attribution-based methods such as integrated gradients or relevance propagation provide more stable estimates of the contribution of individual cells to the learned representations. Furthermore, we found that focusing on the directionality of attention, rather than its magnitude, yields more robust and reproducible interaction signatures across datasets. Complementing this interaction analysis, the linear decoder together with the local and global embeddings enables interpretation of transcriptional programs across spatial scales. Using standardized decoder loadings, we developed a gene evaluation pipeline that identifies scale-specific driver genes in a cell-type-agnostic and hyperparameter-independent manner. More expressive decoder formulations, such as log-sum-exp decoders^53^, may further improve disentanglement of gene programs across embedding dimensions.

Several limitations of the current implementation should be noted. To balance computational feasibility with coverage of large tissue sections, InterScale processes large slides using sliding windows. While this reduces time and memory requirements, it limits the direct interpretation of cellular interactions that span window boundaries. In practice, the diffusion distances of most paracrine signaling molecules are expected to fall within typical window sizes, but long-range endocrine interactions fall outside the scope of the current architecture (Suppl. Fig. 1A). Future work could address this limitation by adopting block-sparse or linear attention variants to input complete tissue slides when analyzing larger spatial datasets^50^. More broadly, even processing the whole slide together is limited to inferring spatial communication within the rough micrometer distance range of the tissue itself, meaning that multi-slide communication or true long-range endocrine signaling across organs remains beyond the scope of this kind of data and approach. Given the lack of ground truth for cell–cell communication, quantitative benchmarking remains challenging. We therefore evaluate InterScale using proxy tasks and consistency with known biological patterns, rather than direct comparison of inferred interactions. Lastly, the interpretation of local and global interaction scales is technology dependent (Suppl. Fig. 1A). For example, bulk-level technologies (e.g. 10X Visum) are restricted to short- and long-range paracrine signalling.

Additional methodological extensions may further improve the framework. InterScale currently represents spatial relationships through direct graph connections between cells. Alternative spatial representations of tissue slides such as graph rewiring^54^ or rotary positional encodings (RoPE)^55^, could improve sensitivity to spatial gradients, density and asymmetric interaction patterns. Additional work could improve the identification of length scales, for example, by adjusting the loss function by weighting local vs global gene contribution based on prior knowledge of the system to be studied or optimizing for their contributions during training. Furthermore, the current framework does not infer directional causal relationships between interacting cell groups. For example, distinguishing whether cell group A influences B independently or via an intermediate group C is beyond the current model’s scope. Finally, discrete sliding windows introduce boundary effects that could be mitigated using adaptive or stochastic window sampling strategies.

Looking forward, InterScale’s framework naturally invites extension to multi-modal or foundation model based architectures^56–59^. The latter could be achieved by adoption of inductive learning frameworks^60^ that would enable InterScale to generalize to previously unseen tissue contexts which may ultimately contribute to virtual cell simulations^61^.

## Methods

### Datasets

#### Dataset 1: Molecular Cartography data

Legnini et al.^37^ studied the sonic hedgehog (SHH) pathway in neurodevelopmental organoids using Molecular Cartography data (Molecular Cartography can be accessed on zenodo https://doi.org/10.5281/zenodo.6143560). They measured the reaction in 4 control and 4 induced organoids. Overall they measured 88 genes with 11.796 control cells from 6 sides and 31.966 SHH induced cells from 11 slides. We preprocessed the data by removing all cells that have zero gene expression and log1p transforming the counts. The authors report that the SHH induction worked the best in organoids 2 and 3 (e.i. slide 4 A2-1 and slide 4 B2-2). We performed the downstream testing on the slides slide4_A2-3 (ctrl) and slide4_B2-2 (SHH) that were excluded from the training and validation set.

#### Dataset 2: CosmX Pancreas

InterScale was trained on a human Type 1 Diabetes (T1D) CosMx datasets from Melton, Jimenez et al., 2025 measuring 386.727 cells and 979 genes (CosMx data can be accessed on 10.5281/zenodo.14870960). The dataset contains 3 slides, each slide with one control and one T1D patient. During preprocessing we log1p-normalized the data, split the data into sliding windows with overlap of 3 and split the slide across train, val and test.

#### Dataset 3: Damond et al, IMC dataset

Damond et al., 2019^62^ collected a human Type 1 Diabetes (T1D) development with IMC (the IMC data can be accessed on https://doi.org/10.17632/cydmwsfztj.1). The dataset consists of 845 images stained with 37 biomarkers from 12 patients with three conditions: recent-onset T1D (<0.5 years, N = 4), long-standing T1D duration (R8 years, N = 4), and controls without diabetes (N = 4). For each donor sections originate from different anatomical regions of the pancreas (tail, body, or head). During preprocessing the raw IMC counts were 99th-percentile-normalized and scaled from 0 to 1 as stated in the publication. We used the squidpy sliding window function with window size 600 and split the patient cases into train (6362, 6278, 6134, 6264, 6180) val (6418, 6126, 6228) and test (6089, 6386, 6380) such that each split represents at least patient from each condition.

#### Dataset 4: Visium, Chen et al

We applied InterScale to the publicly available Chen et al., 2022^30^ data (GSE220442). In the study they collected 10X Visium data from human postmortem brain tissue, three AD patients and three control patients. The data consists of 25.293 spots and 36.501 genes and manually annotated region layers. During preprocessing we removed 245 spots that were manually annotated as noise, performed highly variable genes selection (n=2000) and log1p normalized the gene counts with scanpy. The data was split into train (‘2-3’, ‘18-64’), val (‘2-8’, ‘1-1’) and test set (‘2-5’, ‘T4857’).

### Data loading/preparation

Dataset can consist of single or multiple spatial slides or FOVs with the requirement of equal gene sets.

Each dataset was prepared by normalizing, learning a graph representation using PyTorch Geometric for all cells, if necessary splitting each spatial slide or field of view (FOV) *T* into sliding windows *U*, and then loading the relevant splits as PyTorch Geometric objects into the model for training.

#### Data normalization

Before training, the data input is normalized and log-transformed with scanpy. We observe that InterScales performance works best with maximum normalized values of 4.

#### Graph construction and dataloading

For each sliding window we constructed a graph using the sq.gr.spatial_neighbors from the Squidpy package[R]. Graphs for image-based spatial transcriptomics data are designed by radius based graph to capture density changes across regions and hexagonal grid based graph creation for sequencing-based graphs. The average number of neighborhood connections and standard deviation across radii for the datasets are shown in Suppl. Table 2. To load the graphs into the model we used the Geome package (https://github.com/theislab/geome/tree/main) which creates a PyTorch Geometric object for each sliding window with information about connectivity between nodes and labels. To introduce stochasticity and assess model robustness, we apply a node masking strategy during dataloading. Given a spatial slide or window a percentage *p*_*m*_ of the cells are uniformly sampled. The gene expression value of the cell within the subset of the sampled cell is set to 0.

#### Sliding-window split

We divided each FOV into sliding windows to (1) increase the number of training samples to encourage the model to generalize to cell context in diverse tissue compositions within the same slice, and (2) restrict the number of cells to a feasible sequence length for the transformer for better computational complexity. The sliding window calculation is similar to Kasumi^16^, where each window represents a spatially, rectangularly shaped region. The sliding window function divides a FOV into contiguous, rectangular windows of a given size *w*, with an optional overlap *o* between consecutive windows. If the sliding windows should partially overlap (*o* > 0) then the step size or distance between sliding windows *T* is *T* = *w* − *o*; otherwise the sliding windows are distinct. Note that windows at the border of the size can be smaller than the indicated window size. The sliding window function has been added to SquidPy: https://github.com/scverse/squidpy/blob/main/src/squidpy/tl/_sliding_window.py.

#### Dataset partitions

We randomly split the data into train (70%), validation (20%) and test (10%) set on patient level, such that a data from a patient in the test or validation set has not yet been seen during training. Additionally, the splits are stratified such that each partition should contain an approximately equal amount of data from covariates such as condition.

### Models

We provide a modular coding framework that flexibly allows us to adjust the local and global components of the model. The framework is compatible with Python 3.11 and depends on Scanpy, Squidpy, PyTorch (Lightning), PyTorch Geometric, (optionally WandB) and geome.

- Method: https://github.com/theislab/interscale
- Tutorials: https://github.com/theislab/InterScale_reproducibility/

#### Annotations

Summary of the key annotations used for the model definition:

|  | Notation | Definition |
| --- | --- | --- |
| Dataset | $N \in \mathbb{Z}^+$ | Number of cells/spots/nodes. |
| | $F \in \mathbb{Z}^+$ | Number of genes/features. |
| | $C \in \mathbb{Z}^+$ | Number of prediction classes (e.i. condition, cell type, region) |
| | $P \in [0, 1]$ | Percentage masked nodes, e.i. 0.3 meaning 30% of nodes in input are masked |
| | $B \in \mathbb{Z}^+$ | Number of batches |
| | $T \in \mathbb{Z}^+$ | Number of spatial slides or sliding windows |
| Model input | $X \in \mathbb{R}^{N,F}$ | Gene expression matrix |
| | $A \in \{0, 1\}^{N,N}$ | Adjacency matrix (1: edge, 0: no edge) |
| | $E \in \mathbb{Z}^+$ | Embedding dimension of model. Local and global component dimensions are equal |
| | $H_{local} \in \mathbb{R}^{N,E}$ | Local embedding |
| | $H_{global} \in \mathbb{R}^{N,E}$ | Global embedding |
| | $L \in \mathbb{Z}^+$ | Number of NN layers |
| | $S \in \mathbb{Z}^+$ | Sequence length of transformer/input |
| | $M_{pad} \in \mathbb{R}^{B,S,S}$ | Sequence padding mask |
| | $M_{attn} \in \mathbb{R}^{E, B, S, S}$ | attention mask |
| | $Attn \in \mathbb{R}^{B, S, S}$ | Attention matrix |
| | $D \in \mathbb{Z}^+$ | Number of attention heads |
| | $f_{linear-LD}, f_{linear-GD}$ | Linear Local and global decoder |
| | $G_{Local} \in \mathbb{R}^{F,E},$<br>$G_{Global} \in \mathbb{R}^{F,E}$ | Local and Global gene loadings |

#### Model input

The input to the models are the (1) normalized gene expression matrix *X* ∈ *R*^*N,F*^, where *N* is the number of cells and *F* is the number of genes, and (2) adjacency matrix *A* ∈ [0, 1] (see Graph Construction) and optionally, observed node or graph labels for classification tasks. All models can predict either graph classification, node classification or gene expression regression prediction. In the subsequent sections the different model components used are described.

#### Local component

In this section, we describe the four components implemented in InterScale for the Local Component: two types of graph neural networks: 1) a Graph Convolution Neural Network (GCN) and 2) a Graph Isomorphism Network (GIN), 3) scVI^63^ and 4) precomputed embedding structure. Note that although scVI is included in the Local Component, it primarily captures gene expression structure rather than local spatial information, unlike the other alternatives. For simplicity, we will refer to the local embedding as by *H*_*local*_ ∈ *R*^*N,E*^.

##### Graph Neural Networks (GNN)

We implemented two versions of GNNs: Graph Convolutional Network (GCN) and Graph Isomorphism Network (GIN). Both implementations model the local embedding *H*_*local*_ as a function *f*_*GNN*_ (*X, A*) of the gene expression matrix *X* and adjacency matrix *A*. In both cases, the input layer corresponds to the gene expression values *H*^(0)^ = *X*, which are subsequently updated by information from neighboring nodes. The update mechanism varies between the two models.

##### GCN

The GCN layer is defined as: 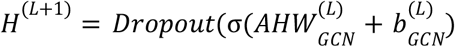 capturing local information by aggregating expression of cells within their spatial neighborhoods using the adjacency matrix representing connectivity. To ensure that each cells expression contributes to the aggregation we incorporated self-connections by adding the identity matrix 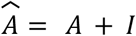. The matrix 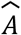 is then row-normalized for equal weighting across neighborhoods, such that each row sums to one. The final output is the GCN embedding *E*_*GCN*_ = *E*_*Local*_. To avoid over-smoothing and -squashing, the GCN layers are fixed to a two-layer depth^23^

##### GIN

The GIN update for node *i* at layer *L* is given by

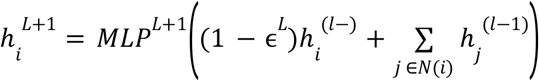

Where *ϵ*^*L*^ is a learnable parameter, *N*_*i*_ is the set of neighbors of node *i*, and *MLP*^*L*+1^ is a Multi-Layer Perceptron. TThe GIN iteratively updates the node embeddings by aggregating information from the neighbors *N*_*i*_ and combining it with the current node representation 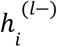, before passing it through the MLP. The final output is the GIN embedding *E*_*GIN*_ = *E*_*Local*_.

##### scVI

scVI^63^ is included as a local component but, unlike the GNNs, it does not explicitly capture spatial neighborhood information. Instead, it learns a low-dimensional latent representation of the gene expression data. We utilize the *scvi*.*nn*.*Encoder* from the scvi-tools package to compute this embedding *E*_*scVI*_, specifically without incorporating batch information. Intuitively, the scVI embedding can be though as replacing the input *X* instead of *E*_*Local*_.

##### Precomputed Embedding

The trainable local component can be replaced by precomputed embeddings, for example from CellCharter^24^ or BANKSY^25^. In this case the locally informed cell representation is computed outside of the InterScale pipeline, saved to the adata.obsm object and retrieved during the training. Precomputed embeddings are not updated and hence a dual decoder architecture (local and global decoder) is not possible.

#### From local to global component

Once we have the local embeddings *H*_*Local*_ ∈ *R*^*N,E*^, we can apply a LayerNormalization to normalize the embedding. Note that compared to GraphTans^22^ we can omit the linear project as we assume equal dimensionality of the local and global embeddings (*E*_*Local*_ = *E*_*Global*_).

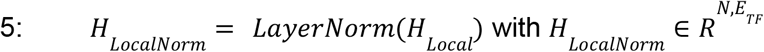

To ensure compatibility with the fixed-length input requirements of the transformer model, we implemented a padding function (Eq. 6). The function *pad_batch* takes as input normalized local node embeddings *H*_*LocalNorm*_ per spatial slide or window to conform to a predefined maximum context length *S*, either by truncating or padding sequences as needed. If the number of cells in a spatial slide or window are larger than the defined context-length (*N* > *S*) then *S* cells are sampled, with priority to masked cells that will be predicted. Otherwise (*N* < *S*) we pad the normalized local embedding with 0s.

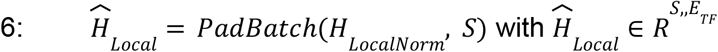

Lastly, a CLS token is added

The output of the transformer is 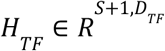 with *S* + 1 because to each sequence (=graph) a CLS token is added, summarizing the state of each graph for graph level prediction.

Details of the steps are described below.

#### Global component

Input: 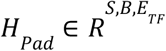

Output: 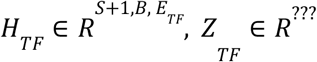

#### Transformer Encoder model

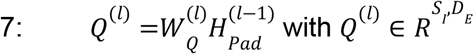

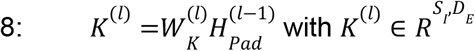

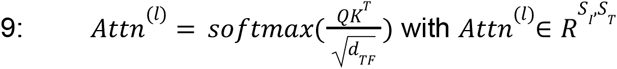

The global component is the TransformerEncoder from the PyTorch implementation following the BERT^64^ architecture. As input, the model receives an embedding representation 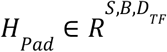 with sequence length *S*, batch size *B* and embedding size *E*_*TF*_. To enforce the transformer to attend to long range cells the attention mask is set to the inverse adjacency matrix mask *M* = 1 − *A*. The mask *M* ∈ *R*^*S, S*^ tells whether two nodes *i* and *j* attend to each other or not. If *M*_*i,j*_ = 0 the nodes are not connected in the local neighborhood (allowing attention) and *M*_*i,j*_ = 0 for locally unconnected nodes (blocking attention). This ensures that the self-attention mechanism in the transformer only considers relationships between nodes that are directly connected in the original local neighborhood. No additional positional embeddings are added to the padded embeddings *H*_*Pad*_ because local information has already been encoded by the GNN and the transformer aims to uncover interactions without distance restrictions.

In PyTorch attn_ouput_weights, with dimensions (B, *S*_*I*_,*S*_*T*_), are calculated according to formula 9 and attn_output according, with dimensions (*S*_*T*_,*E*_*TF*_, *B*), to formula.

Note in the case of self-attention, *S*_*I*_ equals *S*_*T*_.

#### Decoder

For both the local and global components, we have an associated decoder that reconstructs or predicts the required output. The use of separate local and global decoders enforces a clear separation of the learned local and global signals, enabling downstream analyses to attribute inferred effects specifically to either local neighborhood structure or broader tissue-level context.

The output from the respective component, local or global embedding, is then fed into a decoder model. This decoder can predict either graph labels or node labels or the reconstructed gene expression *X*.

Two different types of decoders are implemented: a nonlinear decoder and a linear decoder *f*_*dec*_: 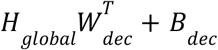 with 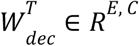. While the nonlinear decoder typically yields superior performance, the linear decoder is directly interpretable and has therefore been used for all reported experiments.

##### Training

The training objective is to minimise the average loss of the local and global decoder output 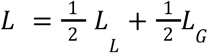. InterScale can learn either 1) graph labels, 2) node labels or 3) reconstructed gene expression. For classification tasks (graph and node label), the Weighted Cross Entropy function 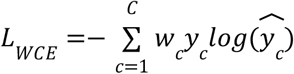 is used. For regression tasks, we use the scaled cosine error loss fun^21^ion19, an adapted version of the cosine error to capture directionality and magnitude of the loss, or maximize the Gaussian negative log-likelihood loss for the masked cells. Loss is calculated on randomly masked cells.

##### Optimization

For optimization we differentiate between parameter space, robustness and hyperparameter tuning. The parameter space optimization addresses the question of: To which extent does an increase in model complexity increase the performance of the model?

##### Hyperparameter tuning

We ran random searches to find the optimal set of hyperparameters for the models and different datasets. For easy reproducibility and model training on new data, we choose the hyperparameters that overall performed best across datasets.

To trade-off performance and runtime we use two different sliding window partitions, meaning that each datapoint is twice represented in the dataset within a different context. Using more sliding window partitions can improve the performance but the user needs to redefine hyperparameters.

##### Robustness

Previous literature has shown that radius and the number of masked nodes to achieve optimal performance. After hyperparameter tuning we test the robustness of the optimally selected hyperparameters for 4 different radius (tissue and dataset specific) and percentage of masked nodes ([0, **0.1, 0.25**, 0.5]) across five seeds (40 - 44).

| Data | Radius |
| --- | --- |
| Legnini et al., 2023 | [0, 100, 200, 300] |
| Melton, Jimenez et al., 2025 | [0, 60, 110, 150] |
| Damon et al., 2019 | [0, 30, 50] |

##### Parameter space sweep

The parameter sweep evaluates the relation between the model performance and trainable model parameters. The goal is to understand whether there is a point of stagnation that further increasing the parameter space leads to no improved model performance. The results are shown in Suppl.Figure 4.

|  |  | Parameter | Values |
| --- | --- | --- | --- |
| Model |  | embed_dim | [16, 32, 64, 128, 256] |
| Local Component | GCN | num_layers | [1,2] |
|  |  | hidden_dim | [32, 64, 128, 256, 512] |
| Global component | Transformer | num_layers | [1,2,4] |
|  |  | n_heads | [2,4] |
|  |  | dim_feedforward | [32, 64, 128, 256, 512] |

### Evaluation

For evaluation we implemented different classification and regression metrics. Classification accuracy of the model was evaluated between predicted and target class based on four different torchmetrics scores: accuracy, F1 micro and F1 marco. The accuracy and F1 micro represent the average across all predictions. Compared to that the macro F1 score provides a balanced measure by accounting for class imbalances.

To evaluate on regression tasks, we evaluated using four different metrics: mean squared error, coefficient of determination (R2 score), pearson correlation and cosine similarity.

#### Inference - Attention relevance

A transformer multi-head self-attention network models the interaction of all cells or neighborhoods across the entire tissue or sliding window. We assume that the attention matrix can be used to assess sender-receiver interactions across cells. One problem is that the interpretation of the attention matrix is not straightforward^65,66^. For example, if different attention layers and heads capture specific data distributions then taking the average will remove those meaningful signals^67^. To obtain a meaningful attention matrix we implement self attention relevance^68,69^ which uses the attention map and gradients to average each head to consider the importance of each head. Finally, the attention map is clamped with a lower bound of zero but no upper bound and all attention scores normalized.

## Downstream tasks

Downstream analysis in InterScale is performed on graph or tissue level and cell level using the attention matrix or on gene level based on the local and global embeddings *H*_*local*_ /*H*_*global*_. A key challenge for attention-matrix-based tasks arises because a large tissue slide is partitioned into smaller, potentially overlapping sliding windows, leading to incomparability across input batches, even within the same slide. This incompatibility is caused by the context-dependent softmax function used to normalize attention scores, which forces them to sum to unity within each specific window. To mitigate this effect for graph-level and cell-level downstream tasks, we apply post-training percentile-based clamping and window-based row-normalization. On the other hand, gene-level evaluation based embeddings remain unaffected by the sliding-window partitions.

### Graph level task: CLS global tissue representation

To aggregate global information for tissue or window-level prediction, InterScale incorporates a *CLS* ∈ *R*^*N*,1^ token into the local embedding. This token acts as a global ‘sink,’ connected to every node (cell) in the global self-attention mechanism, effectively serving as the final, context-aware embedding for the tissue window.

For resistance to extreme transcriptomic outliers or artifacts, the CLS token values are clamped. Specifically, the values are clipped between the 20th (*P*_20_) and 80th (*P*_80_) percentiles calculated across all *CLS* tokens. Let *x*_*i*_ be the original *CLS* token value, *minvalue*_*i*_ = *P*_20_, and *maxvalue*_*i*_ = *P*_80_; the clamped value *y*_*i*_ is computed as*y*_*i*_ = *min*(*max*(*x*_*i*_, *minvalue*_*i*_), *maxvalue*_*i*_). Furthermore, the resulting clamped CLS token vector is scaled by the total number of nodes *N*_*w*_ per window *w* to ensure the aggregated signal is comparable across windows of different sizes.

### Cell level tasks

Cell-level analysis uses the final attention map 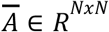 from each sliding window. Each row in 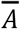 is row-normalized via the softmax function, signifying the relative attention strength received by a cell from all other cells within that window. Direct comparison of attention matrices across different sliding windows is invalid due to the window-specific normalization. To address this and model cellular communication, we focus on extracting local directionality.

To model the directionality of cellular communication across tissue, we compute for each sliding window the net attention flow matrix: 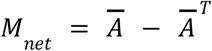. To account for local variations in signal intensity, *M*_*net*_ was normalized 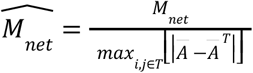 by the maximum absolute net flow within the respective window *T*. In this resulting matrix, the entry 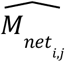 represents the normalized net attention signal flowing from cell *i* to cell *j*.

The net attention matrix 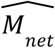 is used for both cell-label net flow aggregation (DotPlot) and for the subsequent spatial flow domain characterization.

### Cell-label net flow interactions

The cell-label net flow is calculated by averaging the cell-label specific net flow. The final positive cell-label net flow is depicted as dotplot (Fig. 4B) such that the size of the dot indicates consistency computed as the reciprocal of the standard deviation and the color the net flow value.

### Spatial Flow Domain Characterization

For each cell *j* the incoming positive net flows were transformed into physical 2D directional vectors. The contribution for each source cell was weighted by a Gaussian distance decay function to prioritize proximal interaction. These single-cell directional vectors were then projected onto a continuous spatial grid using a Gaussian smoothing kernel, generating independent vector fields (*U, V*) for each distinct cell label, used for visualization and identification of spatial flow domains^70^. To characterize the spatial flow landscape, two topological features are extracted for each grid point and cell label: the flow magnitude 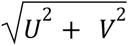 and the vector divergence 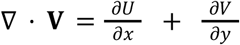. These features are concatenated to form a multi-phenotype “flow signature” for every spatial coordinate. Following feature standardization, we applied the K-means algorithm to identify unsupervised spatial flow domains.

Flow domains were functionally classified based on their average divergence. For each cell type, clusters exhibiting an average divergence significantly above the global mean (μ + *k* σ) were classified as functional “Sources”, while those significantly below (μ − *k* σ) were classified as “Sinks”, with the parameter *k* being user-defined. Finally, to translate these continuous grid annotations back to single-cell resolution, individual cells were assigned to the nearest grid-level flow domain using a KDTree spatial query.

### Gene level tasks

For the gene level analysis we leverage the local and global embedding in addition to the **Linear** Decoder for the local and global component.

### Gene rank analysis

To disentangle which gene interactions the local vs global component captures, we assume that the reconstruction score of the gene expression values inherently capture potential interactions. For both the local and global embedding we learn the reconstructed gene expression matrix 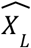 and 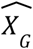 using the local *f*_*linear*−*LD*_ or global linear *f*_*linear*−*GD*_ decoder respectively. The metrics from the evaluation (e.i. coefficient of determination R^2^ or pearson correlation) is applied to quantify how well each gene i is reconstructed given the observed expression values *X* and the predictions 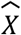. To handle missing values and ensure robust computation, we applied a mask to exclude NaN values from both observed and predicted data, requiring a minimum of two valid data points per gene to compute meaningful metrics scores. For visualization purposes we converted the metric scores into ranks using average ranking to handle tied values, where higher metric scores received higher ranks (indicating better predictive performance).

### Gene Loadings

To interpret the gene-level contributions of the learned latent dimensions, we computed standardized gene loadings separately for the local and global components of InterScale. The local decoder operates on the GCN output, which captures neighborhood-scale structure (*H*_*Local*_ ∈ *R*^*N,E*^), whereas the global decoder operates on the transformer output, which captures broader tissue-scale dependencies (*H*_*Global*_ ∈ *R*^*N,E*^). For the local and global components, predicted expression is given by

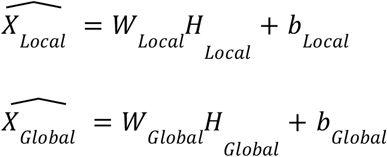

To enable comparison of gene loadings across genes and embedding dimensions, we computed standardized gene loadings analogous to standardized regression coefficients. For each gene *f* and latent dimension *e*, the standardized loading was defined as

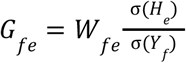

where *W*_*fe*_ is the decoder weight linking latent dimension *e* to gene *f*, σ(*H*_*e*_) is the standard deviation of the corresponding latent dimension across cells, and σ(*X*_*f*_) is the standard deviation of the observed log-normalized expression of gene *f* across cells. This standardization rescales the raw decoder weights to represent the expected change (in standard deviations) of gene expression per one standard deviation change in the latent variable.

### Gene Interpretability pipeline

To interpret the biological content captured by the local and global latent spaces, we implemented a three-step gene interpretability pipeline that progressively moves from latent dimension prioritization to gene selection and, finally, to biological program annotation. This workflow was applied independently to the local and global scales, enabling scale-specific interpretation of the learned representations.

#### 1. Filter embedding dimensions per scale

We first prioritized informative embedding dimensions separately for the local and global components using the standardized gene loading matrices and their corresponding latent embeddings. For a given scale, let *G* ∈ *R*^*F,E*^ denote the matrix of standardized gene loadings, where F is the number of genes and E the number of latent dimensions, and let *H* ∈ *R*^*N,E*^ denote the corresponding latent embedding across N cells.

Dimension importance was quantified using an expression-variance-based score. In the diagonal formulation, the importance of dimension *e* was defined as the squared norm of its standardized loading vector,

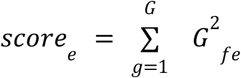

In the full formulation, we additionally accounted for correlations between latent dimensions. Specifically, we computed the latent correlation matrix *Corr*(*H*), the loading cross-product matrix *G*^⊤^*G*, and combined them element-wise. The resulting importance score for each dimension was obtained by summing across rows of the symmetrized matrix,

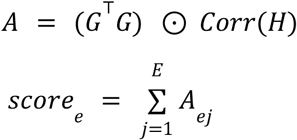

Unless otherwise stated, we used the full mode, as it incorporates both gene-level loading magnitude and correlation structure among latent dimensions. Scores were then normalized to importance ratios by dividing by the total score across dimensions. Dimensions were ranked in decreasing order of importance, and the minimum set of top-ranked dimensions explaining a dataset specific percentage of the cumulative importance was retained for downstream analysis. This procedure was performed independently for the local and global scales, thereby defining scale-specific subsets of informative latent dimensions.

#### 2. Filter genes within selected dimensions

After identifying informative latent dimensions, we selected genes contributing most strongly to each retained dimension. Gene ranking was based on the standardized gene loadings. For each selected dimension, genes were ranked by the absolute value of their loading, such that both positive and negative contributors could be retained as important drivers of the latent axis.

To exclude poorly informative genes, we first applied expression-based filtering using the selected expression layer. Genes were required to be expressed in at least a minimum fraction of cells; optionally, an additional standard deviation threshold could be imposed to remove genes with minimal variability. In some analyses, gene ranking was further weighted by gene-expression variability, using

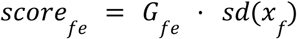

where *G*_*fe*_ is the standardized loading of gene *f* in dimension *e*, and *sd*(*x*_*f*_) is the standard deviation of that gene across cells. This favors genes that are both strongly associated with a latent dimension and transcriptionally variable.

Optionally, we also enforced dimension specificity by requiring each selected gene to contribute more strongly to one retained dimension than to the others. Specificity was quantified either as a ratio or difference between the strongest and second-strongest absolute loading across the selected dimensions. For each retained dimension, the top n genes passing these filters were selected, and the union of these genes across dimensions was used for downstream interpretation. This yielded a compact gene-by-dimension matrix summarizing the dominant transcriptional features captured by each scale-specific latent space.

#### 3. Identify gene programs

To assign biological meaning to the selected gene sets, we performed functional enrichment analysis on the filtered genes using Enrichr through the GSEApy interface. Enrichment was run separately for the gene sets derived from the local and global scales, allowing us to identify biological processes associated with neighborhood-scale versus tissue-scale organization. In this way, the gene interpretability pipeline links scale-specific latent representations to interpretable biological processes, enabling direct comparison of local and global transcriptional programs learned by InterScale.

### Spatial scale-dependent Moran analysis

We quantified spatial autocorrelation across neighborhood scales using Moran’s I on a series of k-nearest-neighbor (kNN) graphs. We used spatial coordinates to build the kNN graph, and the analysis was done either on the learned latent dimensions (local or global) or on the expression of an individual gene. For each feature, we constructed kNN graphs with increasing neighborhood sizes (k = 4, 8, 16, 32, 64) and computed Moran’s I using Squidpy on row-normalized spatial weights. When multiple tissue sections or fields of view were present, graphs were constructed within section using library_key. This yielded a scale-dependent Moran profile I(k) for each latent dimension or gene. We summarized the persistence of spatial structure across scales by computing the area under the Moran curve (AUC) using trapezoidal integration over k. Higher AUC values indicate features with stronger and more persistent spatial organization.

### Benchmarking against competing methods

For comparison to other spatially informed cell-cell communication methods (Suppl. Table 1), we picked two transformer-based methods, multi-scale and spatial transcriptomics data applied methods: AMICI and Steamboat. Both AMICI and Steamboat infer cell-cell communication events from the attention matrix, hence we compared with InterScales attention matrix. Because of the missing ground truths of CCC events we are limited to comparing hypotheses of the methods limiting our benchmark to be ^71^rely similarity based64.

### Methods

**AMICI**^18^ is a single-layer multi-headed cross-attention architecture modelling how receiver cells integrate information from spatial neighbors by generating query embeddings (Q) from learned cell-type-specific parameters that represent receiver identity, while neighboring non-self cell types gene expression profiles generate keys (K) and values (V) through neural network embeddings. Attention scores are computed as distance-weighted softmax over query-key products, where each head learns interaction length scales and sender phenotypes, with the model reconstructing masked receiver expression as a residual from cell-type mean using attention-weighted aggregations of neighbor values. AMICI argues for direct interpretability of pairwise interactions from the attention scores because of the single-layer network architecture. A disadvantage of AMICI is that it relies on cell type annotation which make interaction inference limited to the granularity of the cell type annotation. Additionally, AMICI does not encode the same cell type interactions which simplifies the biological underlying systems.

**Steamboat**^17^ uses a self-attention multi-head attention architecture to decompose gene expression into three spatial scales: within the cell (ego), within a local k-nearest neighborhood (local), and long-range interactions across the whole tissue (global). The cell interactions are learned via a self-supervised mask-predict approach. Each attention head H is assumed to focus on specific interactions and are interpreted separately. Differently to AMICI, does Steamboat not rely on cell type information. However, we noticed that most of Steamboats information is drawn from the ego cell neglecting further reaching interactions.

We compared to four different InterScale implementations:

**Table X:** scVI and PCA are considered as GEX embeddings as they to not represent local neighborhood information.

| Name | GEX embedding | Local Component | Global Component |
| --- | --- | --- | --- |
| GCN |  | GCN | Multi-head<br>Self-Attention<br>Transformer |
| scVI | scVI | - |  |
| PCA | PCA | - |  |
| CellCharter |  | CellCharter |  |

### Legnini et al., 2023: SHH rings

The molecular data from Legnini et al., 2023 does not contain cell types or region annotations of cells. To compare across methods we categorized the cells based on their distance to SHH source. SHH rings are calculated as: First an adaptive threshold for the SHH gene expression is determined using the Otsu method from computer vision (skimage.filters.threshold_otsu). If cells that exceed the defined threshold are labeled as SHH+. Next, the euclidean distance is from every cell to its nearest SHH+ cell is computed using a cKDTree. Finally, the continuous distance values are discretized into four spatially distinct “rings” to defined categorical neighborhoods.

### Correlation of methods

To compare the interaction results from the methods attention matrices we compute the pearson correlation across each attention matrices for all method combinations.. A person correlation of 0 means no correlation, +1 a positive and −1 negative correlation. For the comparison with AMICI, we reset the diagonal of the attention matrix to zero to simulate the equal bias of no self cell type communication.

From our observation inference of interaction for spatial transcriptomics data is challenging for sparse data when the counts per cell are low. Also comparing individual attention scores is difficult as their context and such can vary a lot. Instead we propose to move towards directionality of the attention scores.

## Data availability

All datasets are publicly available.

## Code availability

All models described here are implemented in a Python package available at https://github.com/theislab/interscale. All benchmarking, analysis codes and tutorials are provided at https://github.com/theislab/InterScale_reproducibility/.

## Acknowledgments

We thank Alejandro T., Rushin Grinda, and Mojtaba Bahrami for insightful discussions on the model architecture; Sergio Marco Salas for valuable discussions on applications and downstream analyses; the members of the Theis Lab for helpful discussions; and Robert Gutgesell and Sergio Marco Salas for careful proofreading of the manuscript.

## Author contributions

F.J.T. conceived the study with the help of S.J, F.D and A.C.S.; F.D. and S.J. contributed equally and have the right to list their name first in their curriculum vitae; F.J.T. supervised the project; F.D. and S.J performed the analysis and wrote the code with the help of F.M.; F.D. led the model design and implementation; S.J. led the data analysis. F.D. and S.J. led the data curation with help from T.M.P. and J.B; F.D., S.J and F.J.T wrote the manuscript with the help of F.M. and feedback from N.R.; All authors approved the final manuscript.

