## Supplementary Figures and Methods for "InterScale reveals multi-scale cellular interaction programs in spatial transcriptomics"

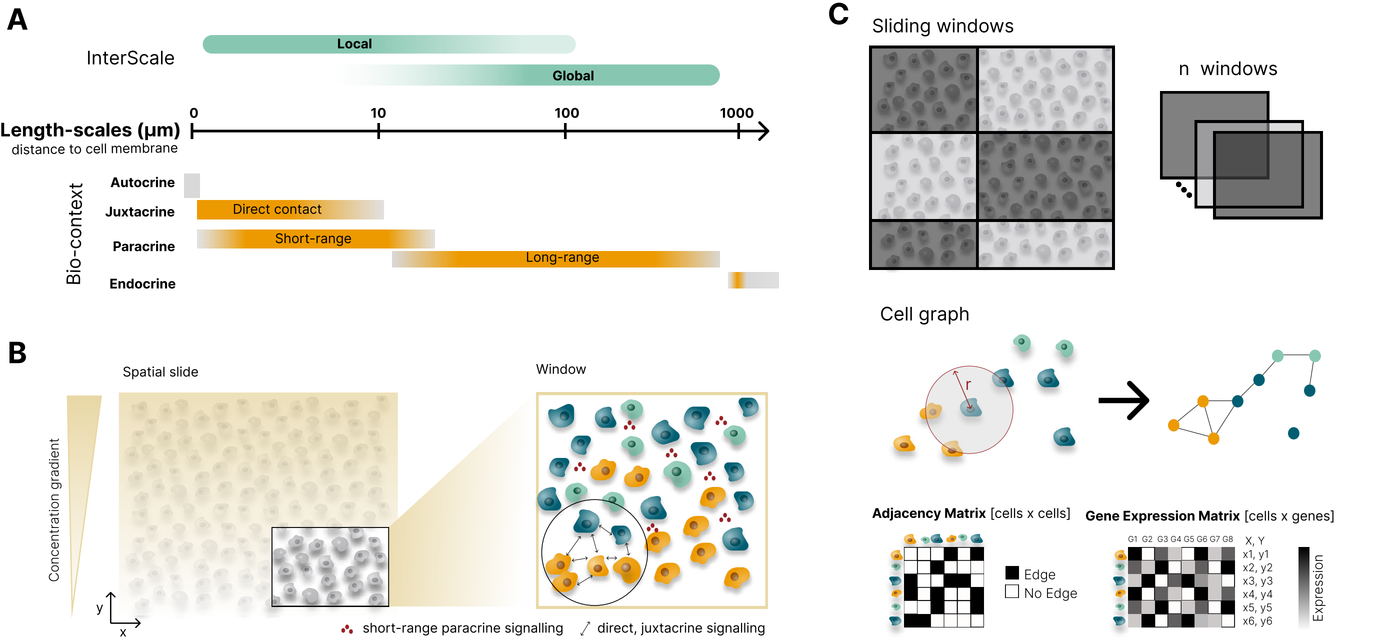
A
InterScale
Local
Global
Length-scales (µm)
distance to cell membrane
0
10
100
1000
Bio-context
Autocrine
Juxtacrine
Direct contact
Paracrine
Short-range
Long-range
Endocrine
B
Spatial slide
Window
Concentration gradient
y
x
short-range paracrine signalling
direct, juxtacrine signalling
C
Sliding windows
n windows
Cell graph
r
Adjacency Matrix [cells x cells]
Edge
No Edge
Gene Expression Matrix [cells x genes]
G1 G2 G3 G4 G5 G6 G7 G8
X, Y
x1, y1
x2, y2
x3, y3
x4, y4
x5, y5
x6, y6
Expression

#### Suppl. Figure 1: Spatial cell-cell communication and data preparation

**A)** InterScale's local and global components are assumed to capture cell-cell communication from different length scales. The local component is more likely to capture interactions up to 10-100µm (autocrine, juxtracrine and short-range paracrine interactions) while the global component is more likely to capture interactions starting from 10-100µm onwards (translating to long-range paracrine signalling). Note that i) the length-scales are represented as gradients because it is a diffusion process without clear boundaries and ii) endocrine signalling across organs is not captured by the global component.

**B)** Illustrates cell-cell interactions on the context of a spatial slide with concentration gradients (such as morphogen gradients) representing long-range paracrine interactions and zooming into a more local view of the spatial slide showing direct contact (arrows) for juxtacrine interactions and short range paracrine signalling.

**C)** The data representation to the model is split into three parts: First, a spatial slide will be split into sliding windows. A cell graph is built on each sliding window or spatial slide, for imaging-based ST that means we connect cells that are within a radius r. The cell graph is represented as an adjacency matrix of size NxN and each node or cell can be associated with a gene expression vector X.

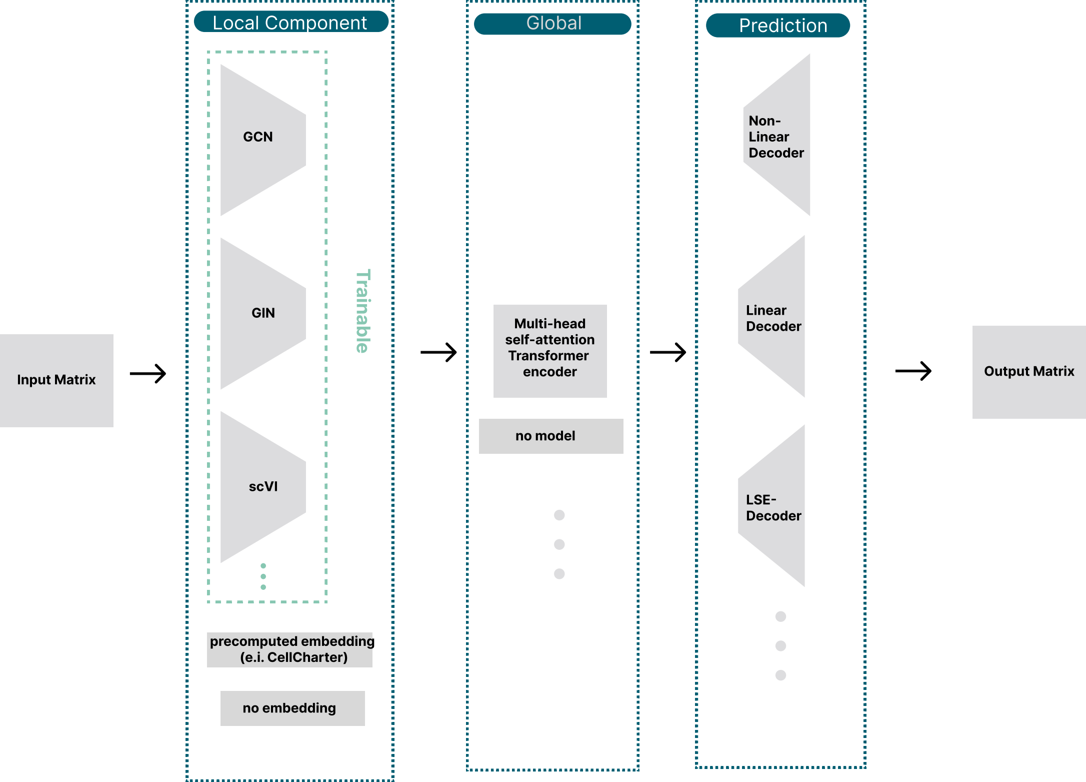
Input Matrix
Local Component
GCN
GIN
scVI
Trainable
precomputed embedding
(e.i. CellCharter)
no embedding
Global
Multi-head
self-attention
Transformer
encoder
no model
Prediction
Non-
Linear
Decoder
Linear
Decoder
LSE-
Decoder
Output Matrix

#### Suppl. Figure 2: Modular Architecture of InterScale

The modular architecture of InterScale consists of three major components: (i) Local Component, (ii) GlobalComponent and (iii) Prediction Component. The input matrix is passed to the (i) Local Component which can either be a trainable module or precomputed embedding. A trainable module can be any type of deep learning model that compresses the gene expression of a node/cell into an embedding space. For example as a graph neural network, (e.i. GCN, GIN) or single-cell specific deep learning model (e.i. scVI). The current implementation of the (ii) global component is a self-attention multi-head transformer with a gradient flow approach to retrieve the attention matrix. For either the local or the global component the user can choose the option of no embedding, which means that the final model represents either local or global structures. Finally, the (iii) prediction components are different types of decoder implementation such as linear and non-linear decoder.

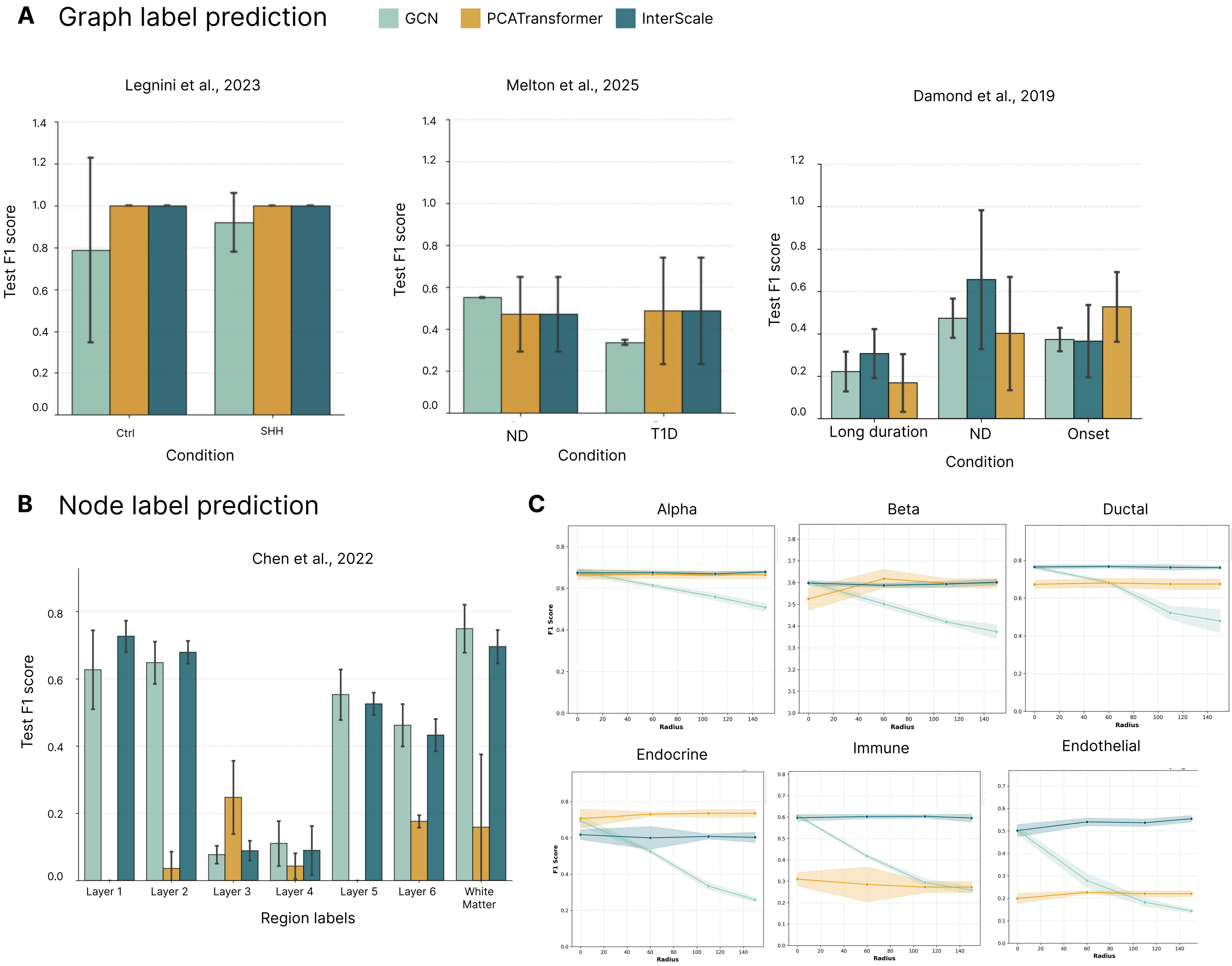

A
Graph label prediction
GCN
PCATransformer
InterScale
Legnini et al., 2023
Test F1 score
Ctrl
SHH
Condition
Melton et al., 2025
Test F1 score
ND
T1D
Condition
Damond et al., 2019
Test F1 score
Long duration
ND
Onset
Condition
B
Node label prediction
Chen et al., 2022
Test F1 score
Layer 1
Layer 2
Layer 3
Layer 4
Layer 5
Layer 6
White Matter
Region labels
C
Alpha
Beta
Ductal
Endocrine
Immune
Endothelial
F1 Score
Radius

#### Suppl Fig. 3: Classification performance of model

**A)** Condition-specific test F1 scores for graph label classification, comparing three model architecture (1) GCN (local component, light blue), (2) PCA-Transformer (global component, yellow), and (3) InterScale (local & global model, dark blue) in their ability to predict the condition associated with a graph. Performance is shown across three distinct spatial omics datasets: molecular cartography neural tube organoid (Legnini et al., 2022, left), CosMx human pancreas (Melton et al., 2025, middle), and Imaging Mass Cytometry human pancreas (Damond et al., 2019, right).

**B-C:** human Alzheimer Disease data

**B)** Top, Bar-plot showing the relative abundance of anatomical layer regions per spatial slide organized by train, validation and test split. Bottom, each line representing a spatial slide showing the aggregated gene count variations per slide colored by condition.

**C)** Test F1 score for node label classification performance of cell types across three model architectures.

**D)** Cell type specific radius sensitivity of model architectures performances for node label prediction on CosMx human pancreas data (Melton et. al, 2025).

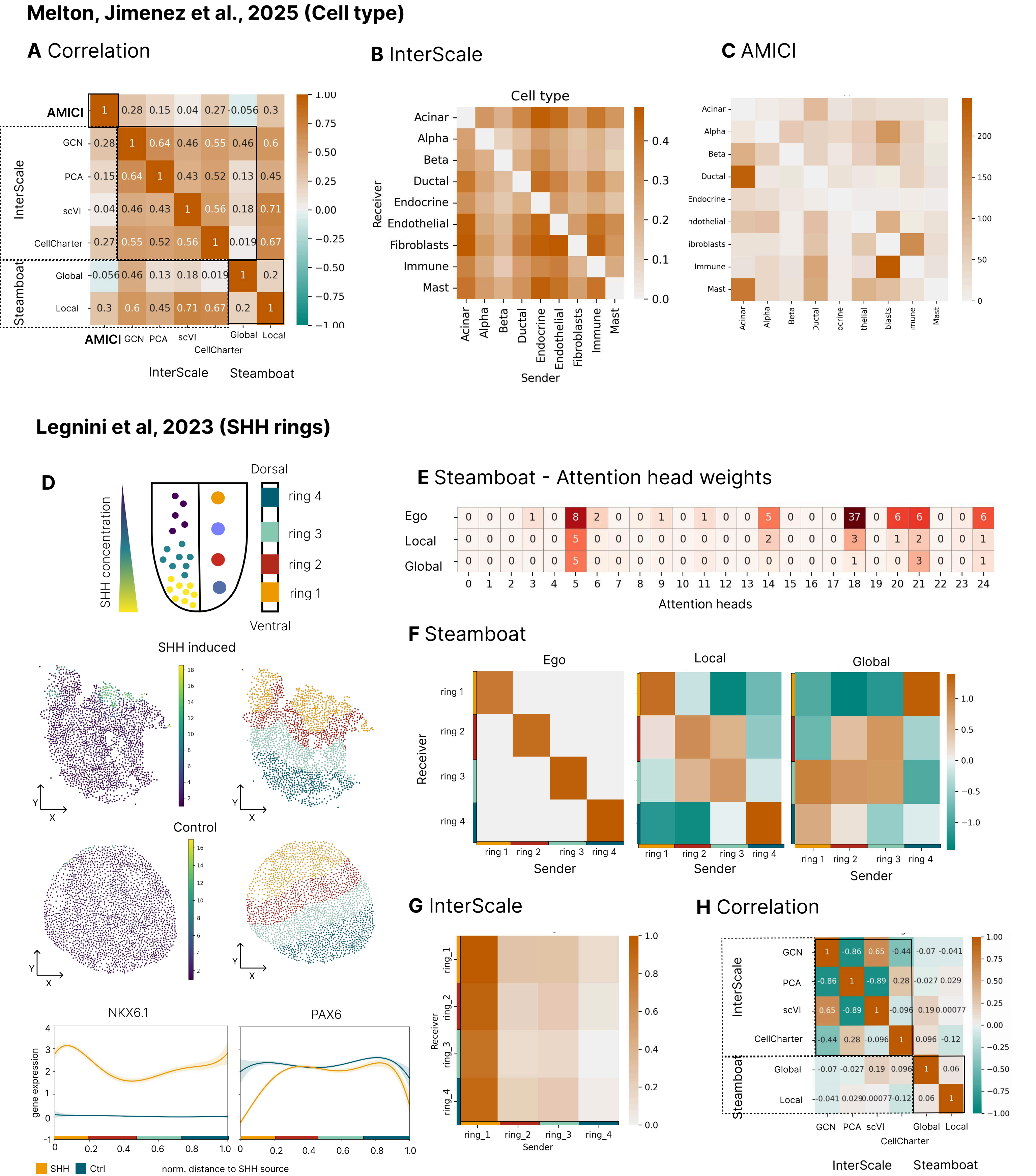

Melton, Jimenez et al., 2025 (Cell type)
A Correlation
B InterScale
C AMICI
AMICI
InterScale
Steamboat
GCN
PCA
scVI
CellCharter
Global
Local
Cell type
Receiver
Sender
Acinar
Alpha
Beta
Ductal
Endocrine
Endothelial
Fibroblasts
Immune
Mast
Legnini et al, 2023 (SHH rings)
D
Dorsal
Ventral
ring 4
ring 3
ring 2
ring 1
SHH concentration
SHH induced
Control
NKX6.1
PAX6
gene expression
norm. distance to SHH source
SHH
Ctrl
E Steamboat - Attention head weights
Ego
Local
Global
Attention heads
F Steamboat
Ego
Local
Global
Receiver
Sender
G InterScale
ring_1
ring_2
ring_3
ring_4
Receiver
Sender
H Correlation
InterScale
Steamboat

#### Suppl. Fig. 4: Benchmarking attention matrix results against AMICI and Steamboat

**A)** Heatmap showing the pairwise Pearson correlation coefficient between aggregated attention matrix from different approaches on Melton, Jimenez et al., 2025 dataset. We tested several input embeddings for the global module of Interscale (GCN embeddings, PCA, scVI embeddings, CellCharter embeddings). We considered both the local and the global aggregated attention matrix from Steamboat and the aggregated attention matrix from AMICI.

B) Heatmap showing the aggregated attention matrix for Melton, Jimenez et al., 2025 dataset produced by Interscale with PCA embedding as input to the Global Component, zeroing the values on the diagonal (same cell type attention) for comparison with AMICI approach.

C) Heatmap showing the aggregated attention matrix for Melton, Jimenez et al., 2025 dataset produced by AMICI.

**D)** To compare attention matrix weights we build distance based rings to the SHH source in control and SHH induced organoid spatial slides (Methods). In increasing distance, ring 1 contains the closest cells to the SHH source and ring 4 the cells furthest away. These rings capture specific gene expression up- and down-regulations that define the dorsal-ventral morphogen gradients in neuronal development.

**E)** Steamboat show the weight of each attention head across three spatial scales: ego, local and global, normalized to a total of 100.

**F)** Aggregated attention matrices for Legnini et al., 2023 dataset produced by Steamboat showing Ego, Local and Global attention scales between SHH rings.

**G)** Heatmap showing the aggregated attention matrix for  Legnini et al., 2023 dataset produced by Interscale with GCN embedding as input to the Global Component between SHH rings.

**H)** Heatmap showing the pairwise Pearson correlation coefficient between aggregated attention matrix from different approaches on Legnini et al., 2023 dataset. We tested several input embeddings for the global module of Interscale (GCN embeddings, PCA, scVI embeddings, CellCharter embeddings). We considered both the local and the global aggregated attention matrix from Steamboat.

The correlation across attention values given the assigned ring category is calculated for C: Steamboat  by spatial scale and D: InterScale's and D: between multiple variants of InterScale and Steamboat.

The correlation of attention values are also compared for Melton, Jimenez et al., 2025 between E: AMICI, Steamboat and multiple of InterScale variants, F: within InterScale attention heads, G: AMICI results.

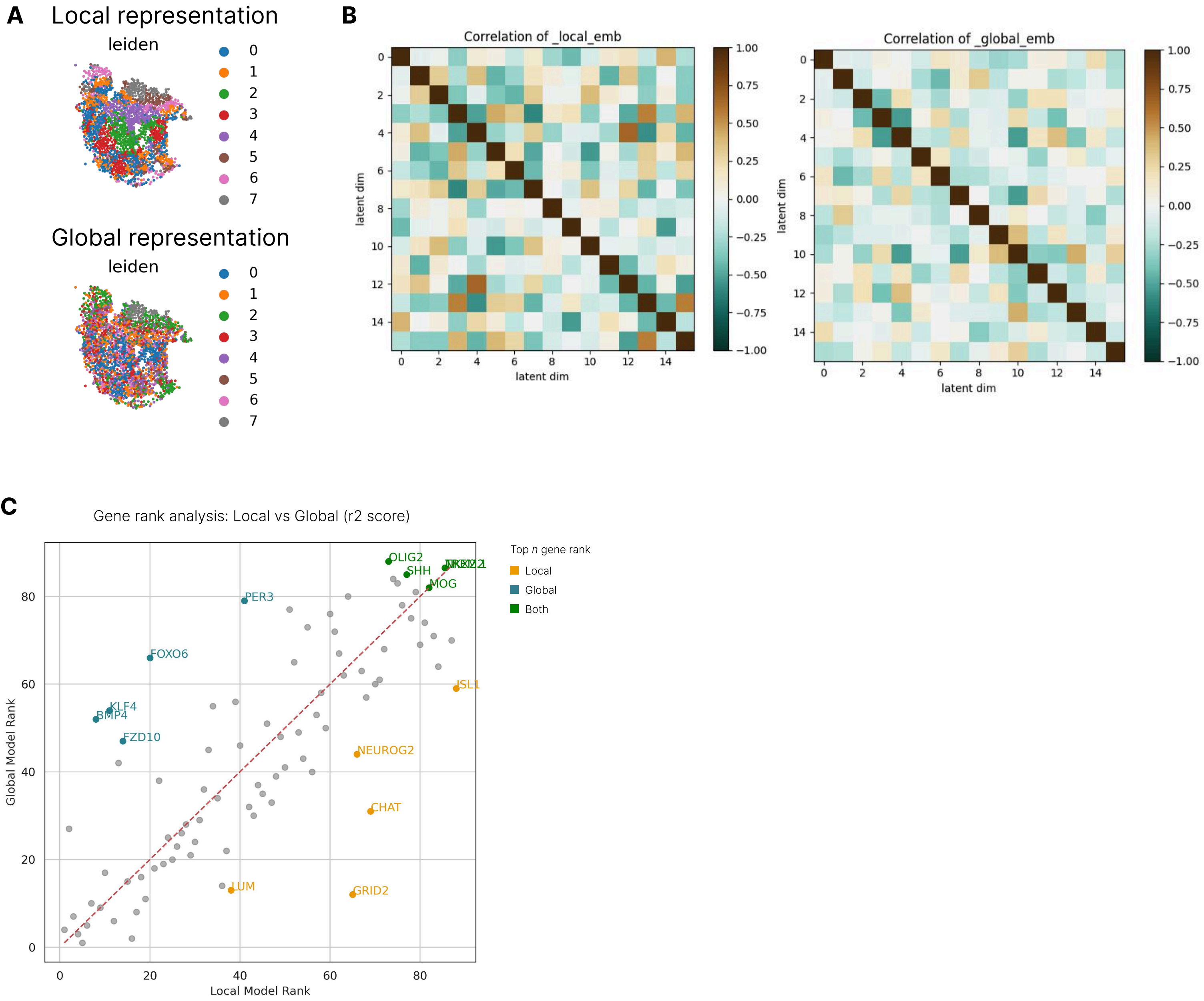
A
Local representation
leiden
0
1
2
3
4
5
6
7
Global representation
leiden
B
Correlation of _local_emb
Correlation of _global_emb
latent dim
latent dim
1.00
0.75
0.50
0.25
0.00
−0.25
−0.50
−0.75
−1.00
C
Gene rank analysis: Local vs Global (r2 score)
Top n gene rank
Local
Global
Both
OLIG2
SHH
MOG
PER3
FOXO6
KLF4
BMP4
FZD10
ISL1
NEUROG2
CHAT
LUM
GRID2
Global Model Rank
Local Model Rank

#### Suppl Fig. 5: Validation of local and global embedding representations in the Legnini et al., 2023 dataset.

**A)** Clustering of cells based on the learned local and global embeddings. Leiden clustering applied to the local embedding space highlights fine-grained transcriptional domains, whereas clustering in the global embedding space captures broader tissue-level organization.

**B)** Pairwise correlation matrices of embedding dimensions for the local (left) and global (right) representations. Low off-diagonal correlations indicate that individual embedding dimensions encode largely independent signals within each spatial scale.

**C)** Gene rank analysis. Each point represents a gene, with the dashed line indicating equal ranking between models. Genes highlighted in orange correspond to those best captured by the local representation, whereas genes highlighted in blue are better explained by the global representation. Genes in green denote those with the highest predictive contribution across models. These comparisons illustrate how local embeddings preferentially capture spatially restricted transcriptional programs, while global embeddings capture broader regulatory signals.

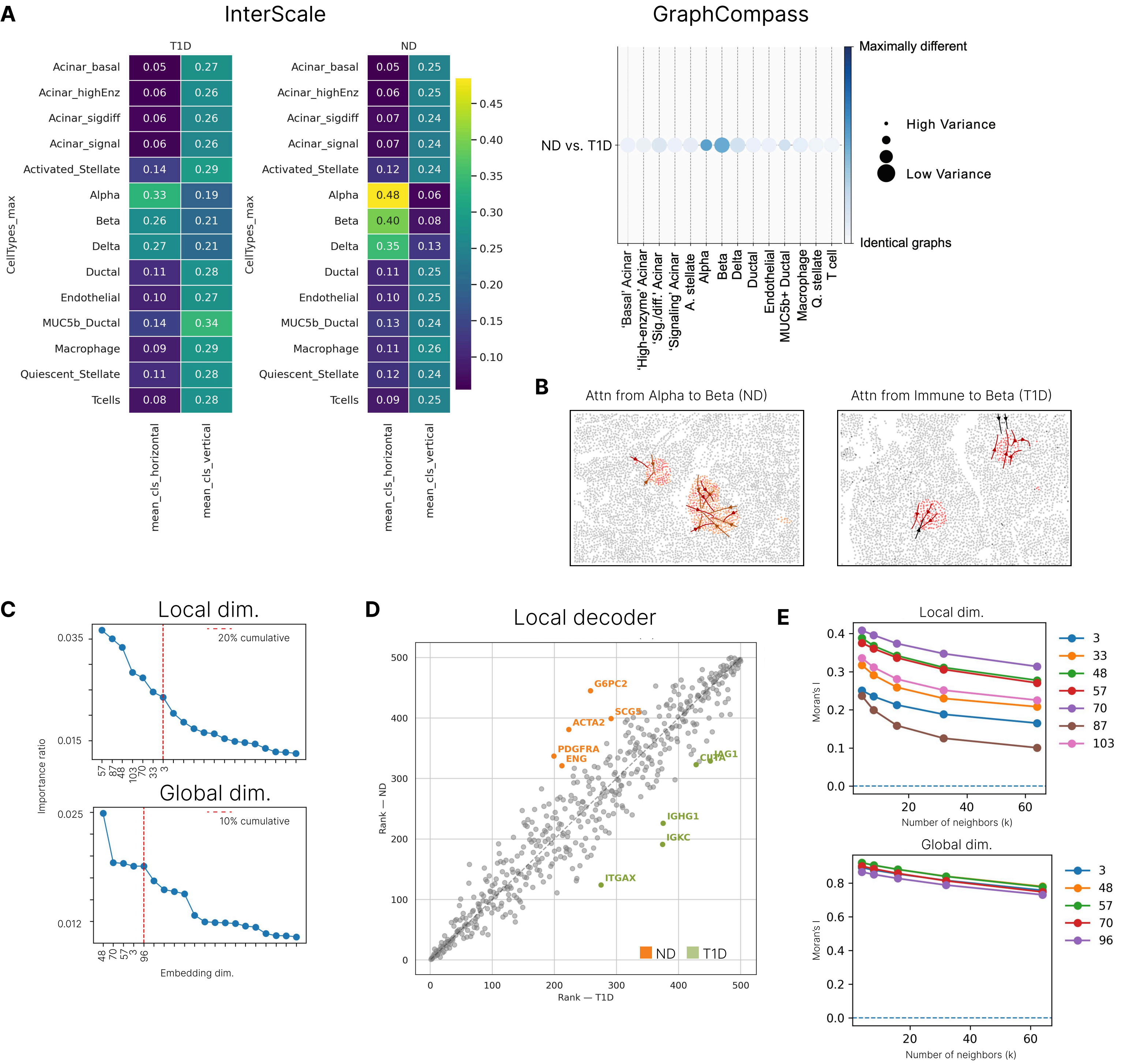

A
InterScale
GraphCompass
T1D
ND
CellTypes_max
Acinar_basal
Acinar_highEnz
Acinar_sigdiff
Acinar_signal
Activated_Stellate
Alpha
Beta
Delta
Ductal
Endothelial
MUC5b_Ductal
Macrophage
Quiescent_Stellate
Tcells
mean_cls_horizontal
mean_cls_vertical
0.05 0.27
0.06 0.26
0.06 0.26
0.06 0.26
0.14 0.29
0.33 0.19
0.26 0.21
0.27 0.21
0.11 0.28
0.10 0.27
0.14 0.34
0.09 0.29
0.11 0.28
0.08 0.28
0.05 0.25
0.06 0.25
0.07 0.24
0.07 0.24
0.12 0.24
0.48 0.06
0.40 0.08
0.35 0.13
0.11 0.25
0.10 0.25
0.13 0.24
0.11 0.26
0.12 0.24
0.09 0.25
0.45
0.40
0.35
0.30
0.25
0.20
0.15
0.10
Maximally different
High Variance
Low Variance
Identical graphs
ND vs. T1D
'Basal' Acinar
'High-enzyme' Acinar
'Sig./diff.' Acinar
'Signaling' Acinar
A. stellate
Alpha
Beta
Delta
Ductal
Endothelial
MUC5b+ Ductal
Macrophage
Q. stellate
T cell
B
Attn from Alpha to Beta (ND)
Attn from Immune to Beta (T1D)
C
Local dim.
20% cumulative
Importance ratio
Global dim.
10% cumulative
Embedding dim.
D
Local decoder
G6PC2
SCG5
ACTA2
PDGFRA
ENG
CUTA
JAG1
IGHG1
IGKC
ITGAX
ND
T1D
Rank — ND
Rank — T1D
E
Local dim.
Global dim.
Moran's I
Number of neighbors (k)

#### Suppl Fig. 6: Comparison of interaction inference, embedding dimensionality, and gene ranking analyses in the human pancreas dataset.

**A)** Comparison between InterScale and GraphCompass interaction analyses for non-diabetic (ND) and type 1 diabetic (T1D) pancreas samples. Left: heatmaps showing average horizontal and vertical CLS attention scores for each cell type inferred by InterScale, representing sender and receiver relevance in tissue-level communication. Right: GraphCompass analysis quantifying graph dissimilarity between ND and T1D conditions across cell types. Color indicates degree of difference between condition-specific graphs, while point size reflects variance across samples.

**B)** Examples of inferred directional interaction patterns visualized on spatial coordinates. Left: attention flow from Alpha to Beta cells in ND samples. Right: attention flow from Immune to Beta cells in T1D samples. Arrows indicate inferred information flow between cells, highlighting localized interaction structures captured by the model.

**C)** Importance ranking of embedding dimensions for local and global representations based on their contribution to standardized gene expression variability. Red dashed lines indicate the cumulative importance thresholds used to select informative embedding dimensions for downstream analyses.

**D)** Gene rank analysis derived from the local decoder between ND and T1D conditions. Each point represents a gene, with the dashed diagonal indicating equal ranking between conditions. Genes highlighted in orange correspond to those more strongly associated with ND-specific programs, whereas green genes show stronger association with T1D-related transcriptional programs.

**E)** Spatial autocorrelation analysis of representative embedding dimensions using Moran's I across increasing neighborhood sizes. Local embedding dimensions exhibit rapid decay of spatial autocorrelation, consistent with spatially confined expression patterns, while global dimensions maintain higher autocorrelation across larger spatial scales.

### Supplements

#### Supplementary Table 1: Overview of Spatial Cell-Cell Communication methods

Suppl. Table 1

We extended the method overview from Armingol et al., 2024[10] that infer cell-cell communication events from spatial omics data. All methods that were already included in the Armingol et al., 2024 overview are marked grey and the newly added methods yellow. Preprints can be identified by the "(PP)" behind the year.

The original columns: Method overview, Additional information, Language, URL, Refs, Year, Real SC, Intracellular, Conditions, Core Function, Deep Learning (DL), Classical Stats, Machine Learning (ML) have been taken over, and appended by Multiscale, Cell types, Spatial representation.

Meaning of the column names:

- Real SC: The model performs inference on single-cell resolution of the data instead of aggregated counts across cell type, neighborhood, etc.
- Intracellular: The model can predict intracellular signalling in addition to LR pairs (e.i. activity of transcription factors and target genes associated with the corresponding receptors).
- Multiscale: Interaction modelling as considered across different length scales. For example, cellular interactions are considered for different neighborhood sizes or different distances to a target/sender cell.
- Cell types: Cell type information are required for either learning or downstream inference.
- Technology: Which spatial technology is the method tested on? (SP = Spatial Proteomics (e.g. IMC), iST: imaging-based Spatial Transcriptomics technologies with single-cell resolution (for example Xenium, CosMx, ...), sST: sequencing-based Spatial Transcriptomics (e.g. Visium))
- Core Functions: Do they use one of the core functions (e.i mean expression between ligand and receptor, correlation) to evaluate the communication score, as shown in Figure 3 in Armingol et al., 2021[1].

### Supplementary methods/notes

#### Data loading

How to load the data for training of the neural network is an important consideration. Some models in spatial transcriptomics, i.e. STELLAR[72], NicheCompass[12] (only 4 neighbors), use the neighborhood data loading approach from GraphSAGE[73]. The NeighbodLoader uniformly samples a fixed-size set of neighbors for N target nodes in the graph. The advantage is the increased efficiency and inductive nature that arguably improves generalization to unseen nodes. There are multiple reasons why we have not used the PyG neighborhood loader for InterScale. First, the NeighborLoader only loads nodes that have a connecting edge to the target node. Not all nodes will be connected because of radius based connections (for delaunay with max cut-off like in CellCharter[24] this wouldn't be a problem, then only remove outliers). Radius based has advantage of representing tissue density. The global component from InterScale should also learn from the node features that are not in the same connected subgraph unit as the target node. Next, if we specify that only a specific number of neighbors are loaded then the node statistics such as node degrees are lost. While this improves the generalizability, the centrality and density could be meaningful in the sense of tissue architecture. For example disease tissues or areas tend to be more densely connected than others. Lastly, the inductive bias in the graph is introduced through masking and selection of nodes in the graph.

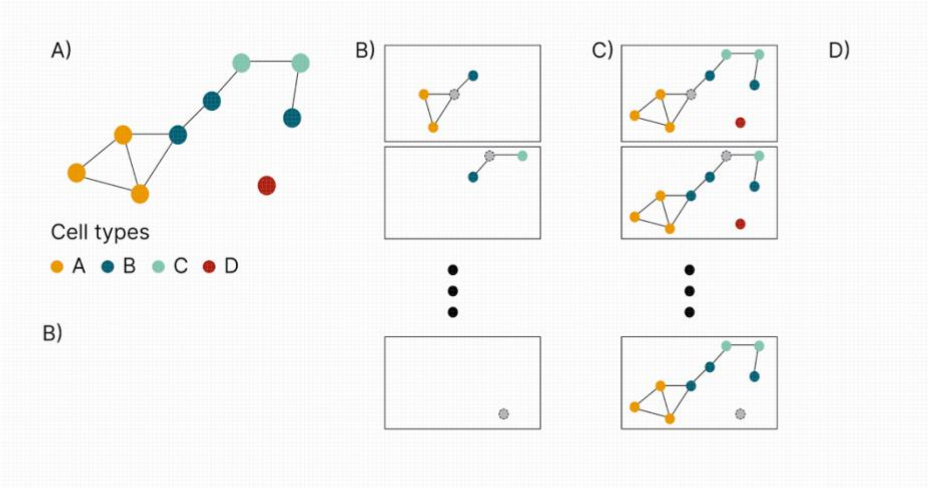


#### Differences to GraphTrans

| | **GraphTrans** | **InterScale** |
|---|---|---|
| **Input** | Feature values, Adjacency matrix | Feature values, Adjacency matrix |
| **Prediction task** | Graph level classification with CLS token | Graph and node-level classification, node-level feature prediction (regression) |
| **Training** | | SSL (masking) |
| **Loss function** | Multi-attribute cross entropy | Classification: (weighted) cross entropy |
| **Transformer attention mask** | No | Attention mask on nodes connected by |
| **Local to global** | If the number of cells N are larger than the max. Sequence length for the transformer, the first [0:S] matrix entries are selected. | If the number of cells N are larger than the max. Sequence length for the transformer then the matrix entries are randomly selected. |

The InterScale architecture is inspired from GraphTrans[22], a graph-transformer architecture that combines a block of GNN, followed by a Transformer. Wu et al., 2021 argue that the node embeddings from the GNN capture the local representations and the transformer calculates their long-range dependencies between each other. One novelty of GraphTrans was the concatenation of a CLS token to the graph embeddings before passing them to the transformer. The performance of GraphTrans was only evaluated for graph-classification tasks utilizing the CLS token. One major finding of the publication was that the classification using the CLS token is more robust than pooling the node embedding outputs of the transformer. As follow up work the authors suggested an extension for GraphTrans was its application to node- and edge-classification.

A limitation of GraphTrans is that, because of the GNN component, over-smoothing and over-squashing remains a concern if the network becomes too deep[74]. For that reason, we limit the GNN depth to two layers. Besides that, if the nodes in the graph are more than the sequence length of the transformer, GraphTrans is disregarding nodes at the end of the input matrix. Hence, it is not considering all node information and instead only focusing on a few. To avoid this inherent bias, in

InterScale we adjusted the local to global transition such that the model randomly samples nodes from the input matrix. Through this step the diversity and information passed to the model is larger.

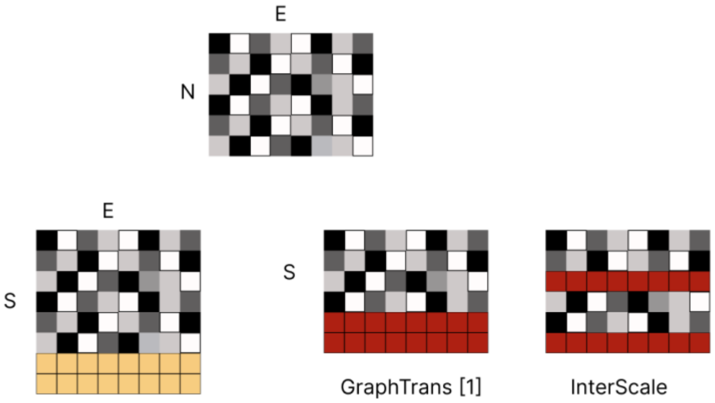


First, in InterScale we move beyond graph classification evaluation from metrics to understanding CLS values within spatial context (plotting them on our spatial slides) and cell information context. Moreover, we extended GraphTrans to node-classification and node-regression. Both tasks rely on the node-embeddings instead of the CLS token, because we are interested in a node-contribution instead of graph-level-wise representation. Lastly, we introduced attention masking of the transformer to hide nodes within the direct neighborhood forcing the model to attend to nodes outside of the local contributions.

The modular architecture implementation is inspired by the GraphGPS framework[74] but we decided for a sequential instead of parallel processing of the input data.

#### Computational complexity of InterScale

The local component used in InterScale is a GCN[23] with time complexity O(Kmd+Knd2) and space complexity O(Knd+Kd2) where n is the total number of nodes, m is the total number of edges, K is the number of layers[75] and d is the number of node hidden features. The global component, is the original multi-head self-attention transformer encoder architecture has O(n2d) complexity per layer L, with sequence length n and embedding dimension[20].

Combining the local and global components, the overall complexity InterScale is O(Knd+Kd2+Ln2d) with the quadratic complexity on the number of cells per tissue slide or sliding window becoming the primary bottleneck.

As mentioned by the authors of GraphTrans[22], the efficiency of the transformer component can be improved by adapting efficient transformer architecture such as LiteTransformer[76] with less FLOPs, Reformer[77] with complexity, and Performer[52] with both less computation and memory complexity.
